# FACT sets a barrier for cell fate reprogramming in *C. elegans* and Human

**DOI:** 10.1101/185116

**Authors:** Ena Kolundzic, Andreas Ofenbauer, Bora Uyar, Anne Sommermeier, Stefanie Seelk, Mei He, Gülkiz Baytek, Altuna Akalin, Sebastian Diecke, Scott A. Lacadie, Baris Tursun

**Affiliations:** Berlin Institute for Medical Systems Biology, Max Delbrück Center for Molecular Medicine in the Helmholtz Association, Berlin, Germany; Department of Biology, Humboldt University, Berlin, Germany; Berlin Institute of Health, Berlin, Germany

## Abstract

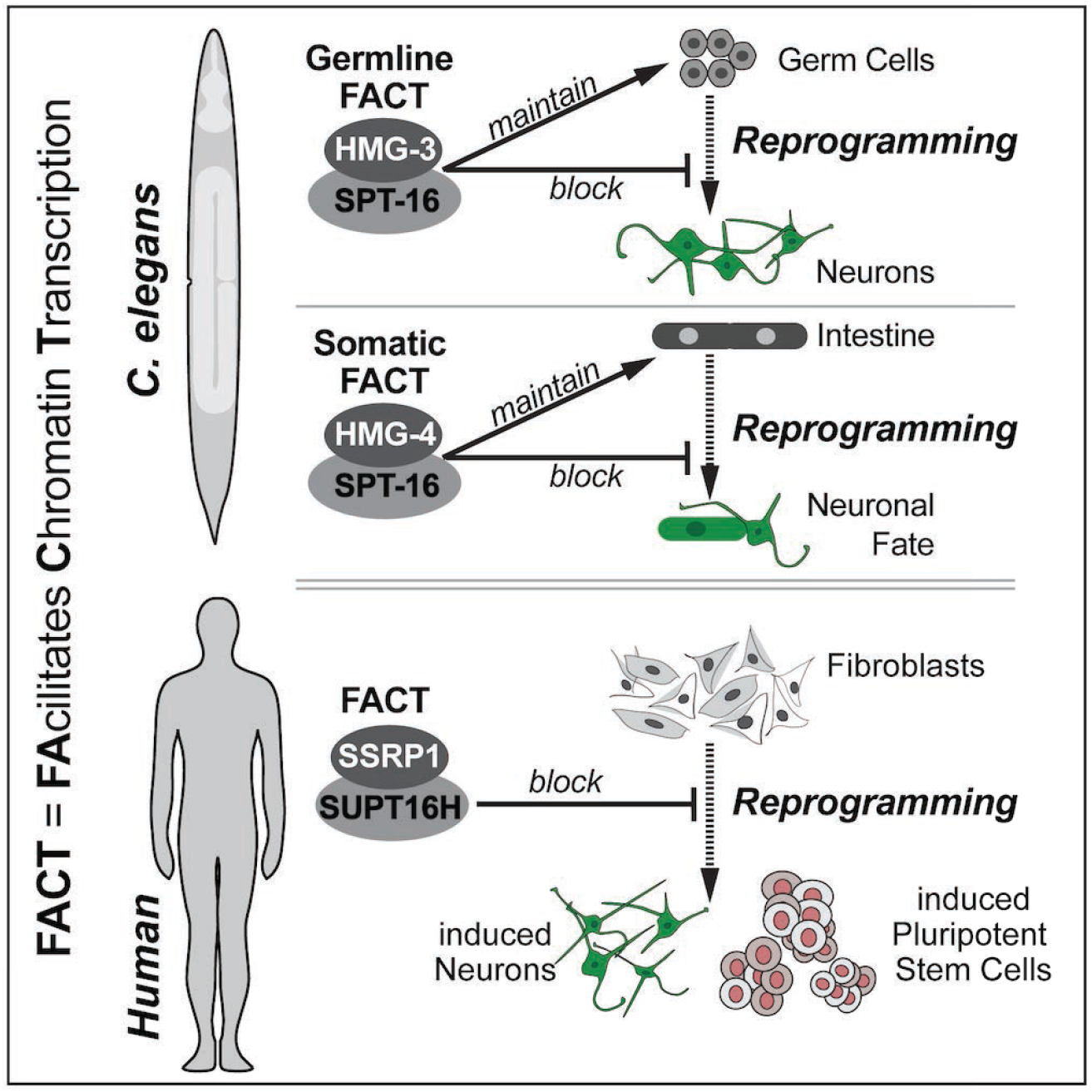

The chromatin regulator FACT (Facilitates Chromatin Transcription) is essential for ensuring stable gene expression by promoting transcription. In a genetic screen using *C. elegans* we identified that FACT maintains cell identities and acts as a barrier for transcription factor-mediated cell fate reprogramming. Strikingly, FACT’s role as a reprogramming barrier is conserved in humans as we show that FACT depletion enhances reprogramming of fibroblasts into stem cells and neurons. Such activity of FACT is unexpected since known reprogramming barriers typically repress gene expression by silencing chromatin. In contrast, FACT is a positive regulator of gene expression suggesting an unprecedented link of cell fate maintenance with counteracting alternative cell identities. This notion is supported by ATAC-seq analysis showing that FACT depletion results in decreased but also increased chromatin accessibility for transcription factors. Our findings identify FACT as a cellular reprogramming barrier in *C. elegans* and humans, revealing an evolutionarily conserved mechanism for cell fate protection.

## Introduction

During development of multicellular organisms cells progressively loose their developmental plasticity and acquire specific fates. Mechanisms that restrict cellular plasticity and safeguard the differentiated state have been in the focus of multiple studies but remain incompletely understood (reviewed by (Guo and Morris, 2017). Both positive as well as negative regulation of gene expression are required during specification and for maintaining cellular identities (Blau and Baltimore, 1991). Positive regulation by transcription factors (TFs) activates genes that define and maintain cellular identities, while negative regulation is important to restrict expression of genes belonging to other cell fates (Blau and Baltimore, 1991). For instance, the human zinc finger protein REST prevents expression of neural genes in non-neuronal cells (Chong et al., 1995) by recruiting epigenetic regulators that silence chromatin (Ballas et al., 2001; Roopra et al., 2004). Notably, such repressive chromatin regulators gained importance in the field of cell fate reprogramming since they can act as barriers for TF-mediated cellular conversion (Becker et al., 2016). Recent examples are the histone chaperones LIN-53 in *C. elegans* (RBBP4/CAF-1P48 in mammals) and CAF-1 in mouse which promote the formation of repressive chromatin and thereby block cell fate reprogramming (Cheloufi et al., 2015; Patel et al., 2012; Tursun et al., 2011). In contrast, chromatin regulators that promote gene expression have not been recognized to act as reprogramming barriers. However, in a genetic screen for factors that safeguard cell fates in *C. elegans* we identified that the histone chaperone FACT (FAcilitates Chromatin Transcription), which is essential for maintaining gene expression (Orphanides et al., 1998; 1999), blocks TF-mediated reprogramming of non-neuronal cells into neurons. FACT is a heterodimer complex consisting of SSRP1 (Structure Specific Recognition Protein 1) and SUPT16H (Suppressor Of Ty 16 Homolog) in mammals (Orphanides et al., 1998; 1999) but has not been studied in *C. elegans* before. We discovered tissue-specific FACT isoforms in the soma and germline, which were not previously described. The somatic FACT is required to maintain the intestinal fate while the germline FACT maintains germ cell identity. The finding that FACT maintains tissue identities and blocks cellular reprogramming implies that positive gene expression regulators contribute to impeding the induction of alternative cell fates. Remarkably, the role of FACT as a reprogramming barrier is evolutionarily conserved as we show that depleting FACT in human fibroblasts significantly enhances reprogramming into induced pluripotent stem cells (iPSCs) or neurons. Our finding points to a deeply conserved mechanism for safeguarding cellular identities revealing FACT as the first positive gene expression regulator that acts as a barrier for reprogramming.

## Results

### The SSRP1 Ortholog HMG-3 Prevents Germ Cell Reprogramming in *C. elegans*

To reveal factors that safeguard cell identities in *C. elegans* we challenged all tissues of the animal with the overexpression of the neuron fate-inducing Zn-finger TF CHE-1, which normally specifies the glutamatergic ASE neuron fate, characterized by expression of the GCY-5 chemoreceptor (Figure 1A and S1A) (Yu et al., 1997). Using a transgenic paradigm that expresses the ASE neuron fate reporter *gcy-5::GFP* and allows ubiquitous CHE-1 expression upon heat-shock (Patel et al., 2012; Tursun et al., 2011) we identified in an F1 RNAi screen that *hmg-3* RNAi allows *gcy-5::GFP* induction in germ cells (Figure 1A). HMG-3 is an ortholog of human SSRP1 (Structure Specific Recognition Protein 1), which, together with SUPT16H (Suppressor Of Ty 16 Homolog), forms the essential chromatin remodeler <u>FA</u>cilitates <u>C</u>hromatin <u>T</u>ranscription (FACT) in humans (Orphanides et al., 1998; 1999). To exclude the possibility that depletion of HMG-3 causes unspecific de-silencing of transgenic reporters we performed *hmg-3^RNAi^* in the absence of CHE-1 overexpression *(che-1^oe^)*. No changes in expression of neither *gcy-5::GFP* (Figure S1B) nor two other reporters that were previously used to detect generic transgene de-silencing (Figure S1D–S1E) (Gaydos et al., 2014; Kelly et al., 2002; Patel and Hobert, 2017) could be observed suggesting that *hmg-3^RNAi^* creates permissiveness for CHE-1 to activate its target gene in germ cells. Induction of *gcy-5::GFP* expression by *che-1^oe^* is accompanied by morphological changes of germ cells showing axo-dendritic-like projections (Figure 1A) indicating that germ cells have converted into neuron-like cells. To assess the extent of germ cell to neuron conversion we examined the nuclear morphology of converted germ cells and expression of other neuronal genes. The *gcy-*5::GFP-positive cells display a nuclear morphology resembling neuronal nuclei (Figure 1B) and expression of pan-neuronal reporter genes such as *rab-3::NLS::RFP* and *unc-119::GFP* (Stefanakis et al., 2015) further demonstrate genuine conversion of germ cells into neuron-like cells in *Iımg-3^mA^’* animals (Figure IB).

**Figure 1.**
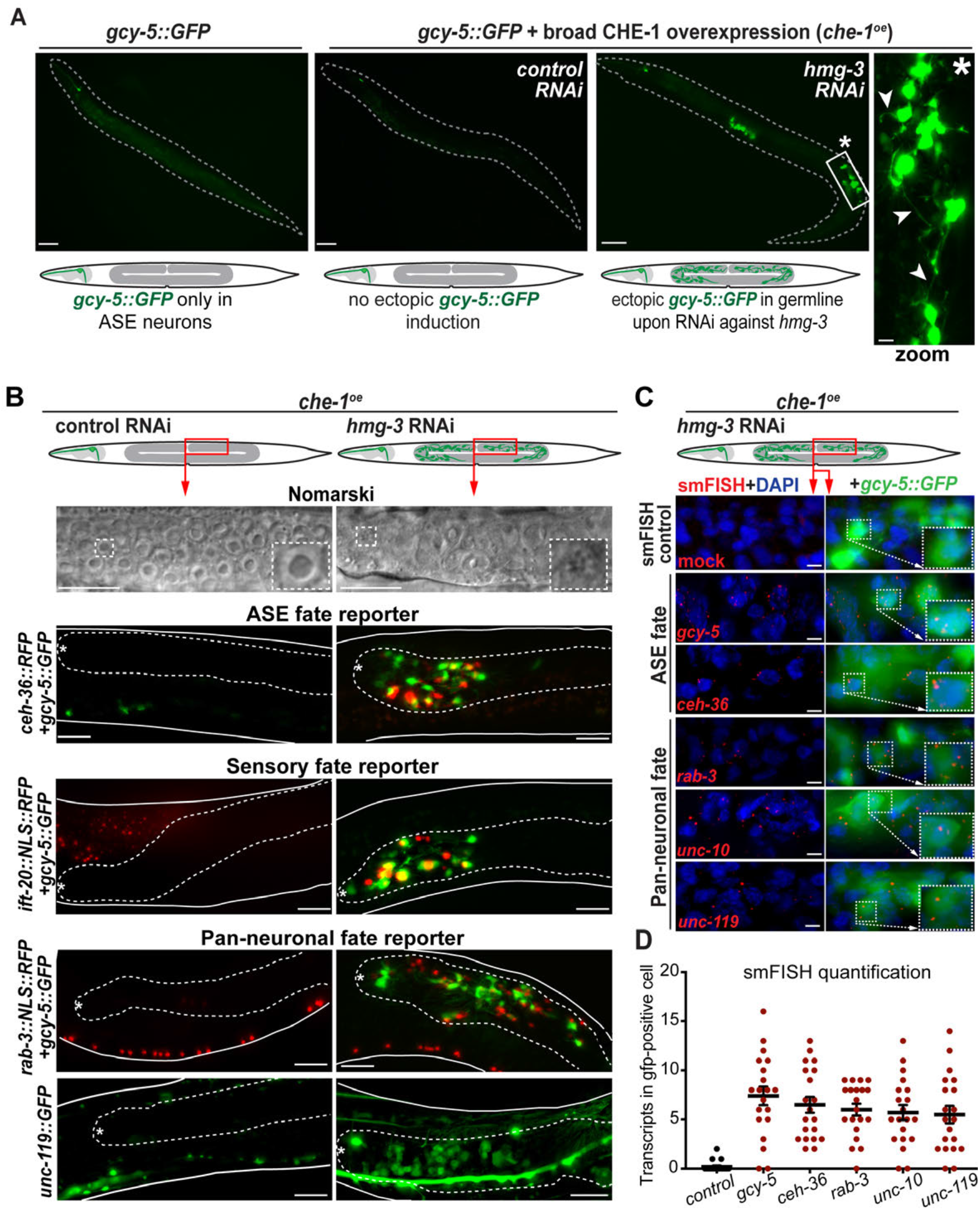
HMG-3, the *C. elegans* Ortholog of SSRP1, Inhibits Reprogramming of Germ Cells to Neurons in *C. elegans*. (A) The glutamatergic ASER neuron is visualized by the ASE fate reporter *gcy-5::GFP*. Broad overexpression of the ASE fate-inducing TF CHE-1 and RNAi against *hmg-3* leads to *gey-5::GFP* induction in germ cells. White arrowheads in the magnified zoom region (white box with asterisk) indicate protrusions resembling axo-dendritic-like projections of neurons in *gcy-5::GFP* positive cells. Dashed lines indicate the outline of animals. For quantification see Figure S1. Scale bars, 20 μm, and 5 μm in the magnification. (B) Differential interference contrast (DIC) microscopy illustrating germ cell nuclei of *hmg-3^RNAt^* + *che-1^oe^* animals with changed nuclear morphology (stippled boxes mark magnification) that resemble neuronal nuclei with speckles. Expression of additional ASE/AWC *(ceh-36)*, sensory *(tft-20)* and pan-neuronal fate markers *(rab-3, unc-119)* can be detected in animals with *gcy-5::GFP* in the germline (outlined by dashed lines). Asterisk labels distal tip of the germline. For quantification see Figure S1. Scale bars, 10 μm. (C) Single molecule fluorescent in situ hybridization (smFISH) to detect transcripts derived from endogenous neuronal genes *gcy-5, ceh-36, rab-3* (Rab3), *unc-10* (RIM), and *uncll9* (UNC119) in *hmg-3^RNAt^* germ cells. mRNA molecules are visualized as red dots. Controls were treated with mock hybridizations. Dashed boxes indicated the magnified area. smFISH probes used in the study are described in STAR methods. Scale bars, 2 μm. (D) Quantification of smFISH based on counts of hybridization signals (red dots) per GFP-positive cells. For each condition, 20 GFP-positive cells were counted for smFISH-derived transcript detection based on fluorescence signals.

Moreover, reprogrammed germ cells also express *tft-20::NLS::RFP*, a marker for ciliated neurons such as ASE, and the ASE/AWC-specific gene *ceh-36::GFP* (Hobert, 2010; 2013) (Figure 1B and S1F). Importantly, transgene reporter expression reflects endogenous expression of neuronal genes as shown by single-molecule FISH (smFISH) to detect neuronal gene transcripts. Transcripts from *gcy-5, ceh-36, rab-3, unc-119*, as well as from the conserved synaptic protein-encoding gene *unc-10* (RIM), become expressed in the reprogrammed germ cells (Figure 1C). Furthermore, the acquisition of neuronal characteristics is accompanied by the loss of germline marker *pie-1* and germ cell morphology (Figure S1G), corroborating the notion that germ cells convert into ASE neuron-like cells in *hmg-3^RNAt^* animals upon induction of CHE-1 expression.

### Specificity of Germ Cell to Neuron Reprogramming in HMG-3 Depleted Animals

To examine whether CHE-1 reprograms germ cells in the *hmg-3^KNAl^* germline to properly specified ASE neurons we tested the expression of markers belonging to other neuron subtypes. CHE-1 does not induce GABAergic or cholinergic neuron reporters in *hmg-3^mAt^* animals (Figures 2A and 2B) arguing that reprogrammed germ cells are not generally mis-specified but acquire a specific glutamatergic ASE neuron fate. We next asked whether *hmg-3* plays a widespread role in preventing germ cell conversion under ectopic expression of TFs. Mis-expression of the GABAergic neuron fate-inducing homeodomain TF UNC-30 (Jin et al., 1994) resulted in germ cell to GABAergic neuron conversion in *ìınıg-3^mAì^* animals (Figures 2C and 2D).

**Figure 2.**
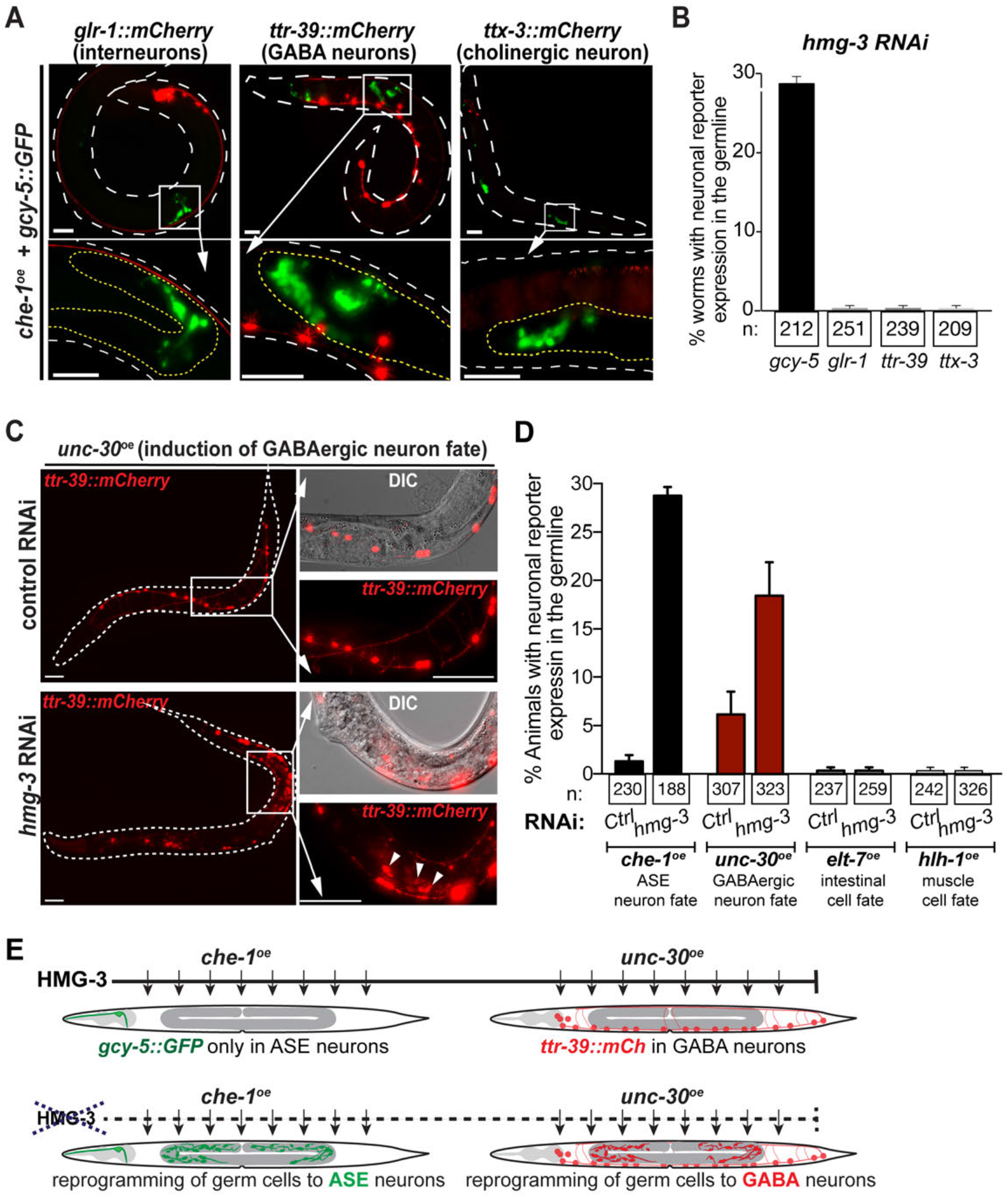
HMG-3 Depletion Allows Conversion of Germ Cells to Specific Neurons. (A and B) Assessment of non-ASE neuron markers after *che-1* “-induced germ cell reprogramming in HMG-3-depleted animals. (A) Representative images and (B) quantification show normal expression of non-ASE markers *gìr-1 ::mCherry* (interneurons), *ttr-39::mCherry* (GABAergic neurons) and *ttx-3::mCherry* (cholinergic interneuron) but not in reprogrammed germ cells (magnifications) expressing *gcy-5::GFP*. Dashed white lines outline worm, yellow lines outline germline. Scale bars, 20, and 5 μm in magnifications. Error bars represent SEM. (C) Representative images for induction of the GABA fate reporter *ttr-39::mCherry* in the germ line of *hmg-3^RNAt^* animals upon overexpression of GABA neuron-fate inducing TF UNC-30 *(unc-30^oe^)*. Dashed lines indicated the outline of the worm, boxes indicate the magnification area and arrows in the zoom indicate reprogrammed germ cells. Scale bars, 20 mm, and 5 mM in magnifications. (D) Quantification of germ cell conversion upon induction of different fate-inducing TFs. Neuronal fate induction can be detected in 20-30% animals after *che-1^oe^* (ASE neuron) or *unc-30^oe^* (GABA neuron), but no intestinal or muscle fate induction by over-expressed ELT-7 *(elt-7^oe^)* or HLH-1 *(hlh-1^oe^)*, respectively. Error bars represent SEM. (E) Schematic representation of transgenic animals for ASE neuron fate induced by zinc-finger TF CHE-1 and GABA neuron fate induced by *unc-30P^e^. che-1^oe^* and *unc-30o^e^* in adults after hmg-3 RNAi induces reprogramming of germ cell to ASE and GABA neurons, respectively. However, mis-expression of the myogenic bHLH TF HLH-1 (MyoD homolog) (Harfe et al., 1998) or the intestinal-fate inducing GATA-type TF ELT-7 (Riddle et al., 2013) failed to convert germ cells into muscle or gut-like cells, respectively (Figures 2D and S2A-B).

This suggests that *hmg-3^mAl^* specifically creates permissiveness for germ cell to neuron reprogramming. Furthermore, we tested whether mitotic or meiotic germ cells are susceptible to being reprogrammed into neurons. We used animals that carry a temperature-sensitive gain-of-function mutation of the Notch receptor GLP-1 that causes loss of meiotic germ cells (Pepper et al., 2003). Growth at the non-permissive temperature would lead to loss of germ cell to neuron reprogramming if the converting germ cells belong to the meiotic pool. However, germ cell conversion is not lost in the *glp-1(gf)* mutant background, suggesting that mitotic rather than meiotic germ cells are the source for reprogramming into neuon-like cells (Figure S2C). This reprogramming is also independent of the cell cycle activity of germ cells, as blocking cell cycle progression by Hydroxyurea (HU) (Gartner et al., 2004) did not inhibit germ cell to neuron reprogramming in *hmg-3^mAl^* animals (Figure S2D). Taken together, depletion of the SSRP1 ortholog HMG-3 in *C. elegans* allows reprogramming of mitotic germ cells into specific neurons upon expression of a neuron fate-inducing TFs.

### Other FACT Subunits Prevent Neuron Fate Induction in the Intestine

RNAi in embryos against a number of genes causes lethality during early development which was also previously observed (Kamath et al., 2003). Therefore, we additionally performed RNAi only after birth of the animals and found that depletion of another SSRP1 homolog, *hmg-4* as well as a previously uncharacterized *C. elegans* gene *F55A3.3* allows highly penetrant ectopic *gcy-5::GFP* induction in intestinal cells (Figures 3A-3B). Strikingly, *F55A3.3* encodes for the sole *C. elegans* homolog of the human FACT subunit SUPT16H (Guindon et al., 2010; Ruan et al., 2008). We therefore refer to *F55A3.3* as *spt-16* hereafter. While reprogrammed germ cells in *hmg-3^mAl^* animals display neuronal morphology, the intestinal cells with *gcy-5::GFP* in *hmg-4^mAl^* or *spt-16^mAl^* maintain their original morphological features indicating an incomplete cell conversion (Figure 3A). However, smFISH revealed that the *gcy-5::(ĵl.’ľ-posìú\c* gut cells show expression of neuronal genes as seen in reprogrammed germ cells (Figures 3C-3E).

**Figure 3.**
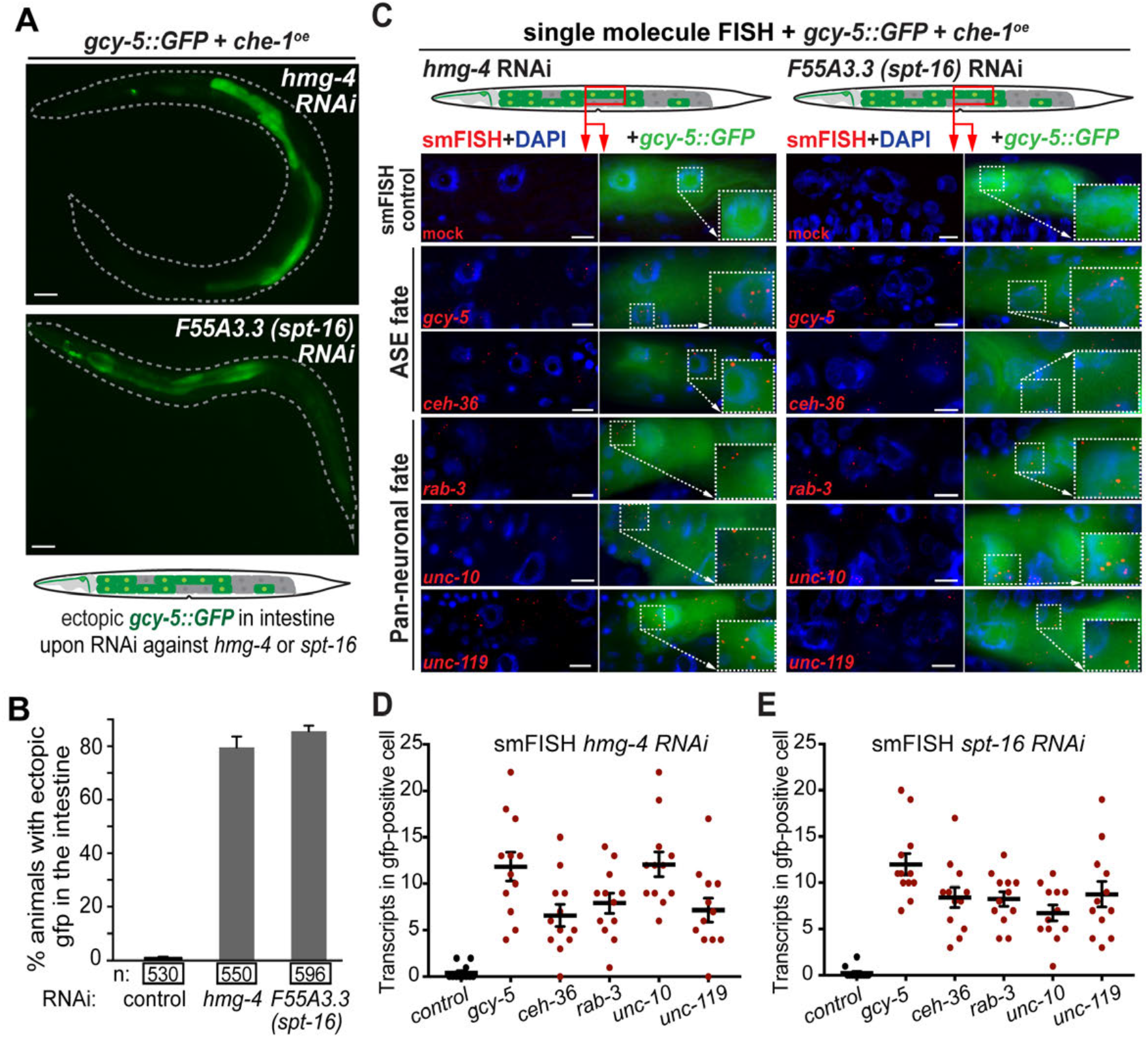
Other FACT Subunits Prevent Neuron Fate Induction in the Intestine. (A) RNAi after embryonic development against *hmg-4*, encoding for another SSRP1 ortholog, and the uncharacterized gene *F55A3.3*, encoding the SUPTH16 ortholog and therefore termed *spt-16* in our study, allows *gcy-5::GFP* induction in the intestine. Scale bars, 20 μm. (B) Quantification of animals showing *gcy-5::GFP* in the intestine in *che-1^oe^* + *hmg-4^RNAt^* or *spt-16^RNAt^* animals. More than 80% of *hmg-4^RNAt^* and *spt-16^RNAl^* animals show ectopic *gcy-5::GFP* induction in the gut cells. Error bars represent SEM. (C) Detection of endogenous transcripts derived from neuronal genes in *hmg-4^RNAt^* and *spt-16^RNAt^* intestinal cells using smFISH as described before in Figure 1C. Individual mRNA molecules are visualized as red dots. Controls were treated with mock hybridizations. Dashed boxes indicate the magnified area. smFISH probes used in the study are described in STAR methods. Scale bars, 2 μm. (D and E) Quantification of smFISH based on counts of hybridization signals (red dots) per GFP-positive cells. (D) Quantification of neuronal transcripts in the intestine upon *hmg-4* RNAi. (E) Quantification of neuronal transcripts in the intestine upon *spt-16* RNAi. For each condition, 20 GFP-positive cells were counted for smFISH-derived transcript detection based on fluorescence signals as exemplified in (C).

The lack of morphological changes towards a neuron-like appearance might be due to the intestine being structurally more specialized and constrained than germ cells. Nevertheless, intestinal cells switched to a neuronal gene expression profile, which is stably maintained even 2 days after CHE-1 induction similarly to HMG-3 depletion mediated germ cell to neuron reprogramming (Figures S2E and S2F). Overall, HMG-4 and SPT-16 are preventing the induction of neuronal genes in the intestine and identification of all *C. elegans* FACT subunits in our screen indicates that FACT plays a general role in safeguarding cellular identities in different tissues.

### *C. elegans* has Germline and Soma-specific FACT Isoforms

The tissue-specific effects of depleting different FACT subunits on ectopic *gcy-5::GFP* induction suggested distinct expression patterns of the FACT genes. HMG-3 and HMG-4 both share more than 90% amino acid homology with SSRP1 (Figure 4A) (wormbase.org) and with each other. Both cross-react with polyclonal anti-HMG-3/4 antibodies labeling all cells of the animal when used for immunostaining (Figure 4B). To discriminate between HMG-3 and HMG-4 we tagged HMG-3 with the HA epitope using CRISPR/Cas9 editing. Anti-HA antibody immunostaining revealed that HMG-3 is exclusively expressed in the germline explaining the distinct effect of *hmg-3^mAl^* on permissiveness for reprogramming germ cells (Figures 1B and 4C). In contrast, a fosmid-based fluorescent reporter for HMG-4 revealed exclusive soma expression with high intensity in the intestine (Figures 4D and S2G) while SPT-16 expression could be detected in all tissues, also with predominance in the intestine (Figure 4E).

**Figure 4.**
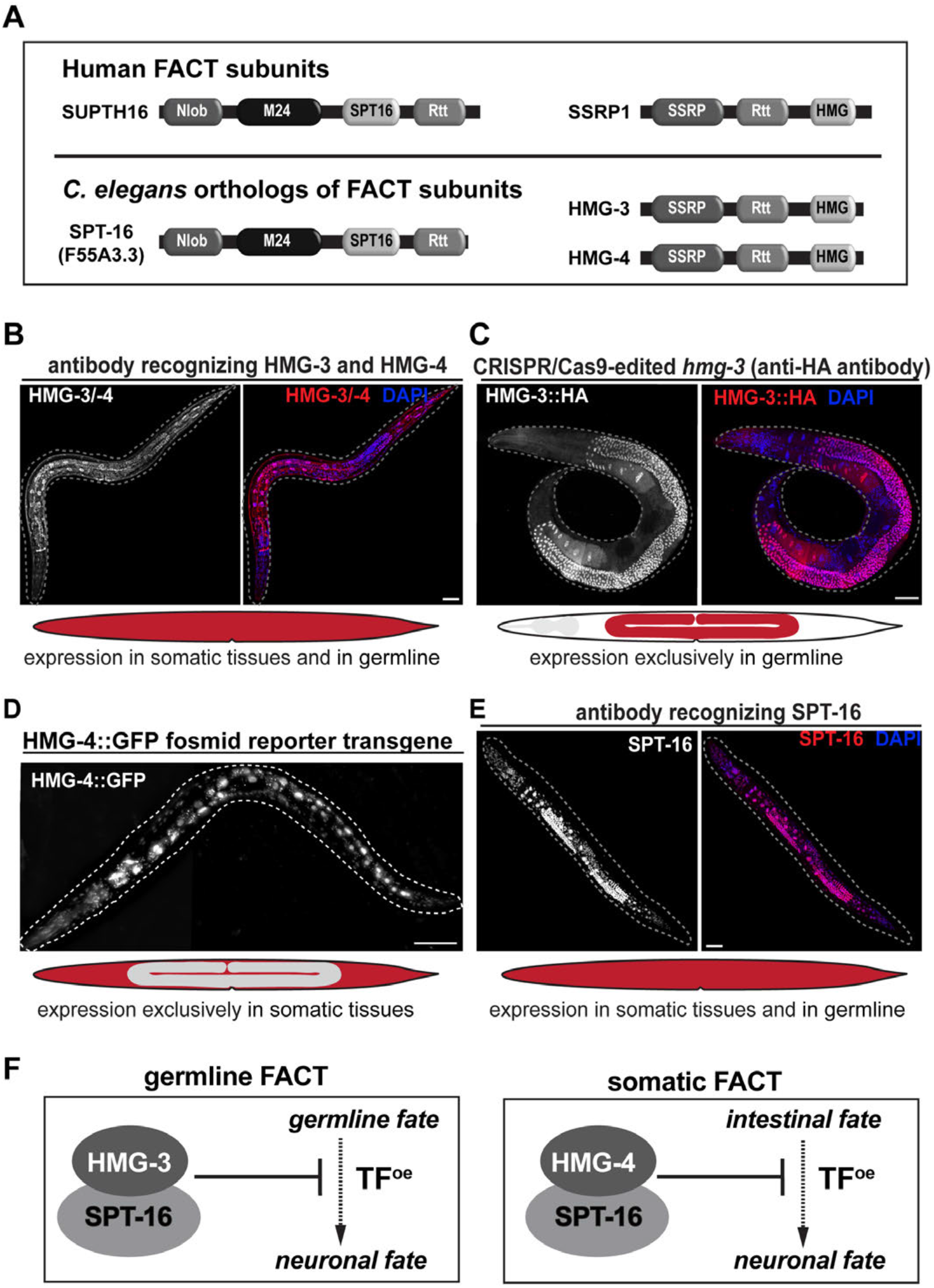
C. *elegans* Has Soma-Specific and Germline-Specific FACT Isoforms. (A) Models of FACT subunits in Human and *C. elegans*. Conserved protein domains according to *Pfam* (pfam.xfam.org) and *InterPro* (ebi.ac.uk/interpro) are indicated: Nlob = N-terminal lobe domain in FACT complex subunit Spt16; M24 = Metallopeptidase family M24; SPT16 = FACT complex subunit Spt16p/Cdc68p; Rtt = Histone chaperone Rttp106-like; SSRP1 = structure-specific recognition protein; HMG = High mobility group box domain. (B-E) Immunostaining of FACT subunits in L4 stage worms. (B) The highly similar HMG-3 and HMG-4 proteins cross react with antibodies and display expression in the whole adult worm. (C) Immunostaining of CRISPR-tagged *hmg-3* with HA shows that HMG-3 is expressed exclusively in the germline. (D) A fosmid-based *hmg-4::GFP* reporter reveals that HMG-4 is expressed in the intestine and other somatic tissues but not in the germline (Figure S2G). (E) Specific antibodies against SPT-16 reveal expression in somatic tissues with high levels in the intestine as well as in the germline. Dashed lines indicate outline of the animals. Scale bars, 20 μm. (F) Expression patterns and phenotypes suggest germline-specific FACT is formed by HMG-3 and SPT-16 and somatic FACT is composed of HMG-4 and SPT-16.

Though *spt-16* is expressed in the germline we could not assay for conversion of germ cells to neurons in *spt-16* RNAi animals. Reprogramming of germ cells requires F1 RNAi as shown for the depletion of *hmg-3* or other previously identified factors (Patel et al., 2012; Tursun et al., 2011). However, exposure to *spt-16* RNAi, and also *hmg-4* RNAi, during embryonic development caused early lethality such that RNAi could only be applied post-embryonically. Nevertheless, the specific RNAi effects together with the mutually exclusive expression patterns of *hmg-3* and *hmg-4* suggest that HMG-3 and SPT-16 form a germline-specific FACT that safeguards germ cells while HMG-4 and SPT-16 constitute the somatic FACT protecting the intestinal fate (Figure 4F) in *C. elegans*.

### FACT Depletion Affects Chromatin Accessibility of TFs and Cell Fate Maintenance

FACT promotes gene expression by nucleosome dis-assembly and assembly (reviewed in (Hammond et al., 2017; Reddy et al., 2017)), suggesting that loss of FACT causes alterations in chromatin. We therefore profiled changes in chromatin accessibility using the Assay for Transposase-Accessible Chromatin with high-throughput sequencing (ATAC-seq) (Buenrostro et al., 2013). ATAC-seq was performed with FACT-depleted whole worms without inducing *che-1* overexpression in order to detect chromatin alterations that were caused by FACT depletion but not due to the activity of the reprogramming TF (Figure 5A).

**Figure 5.**
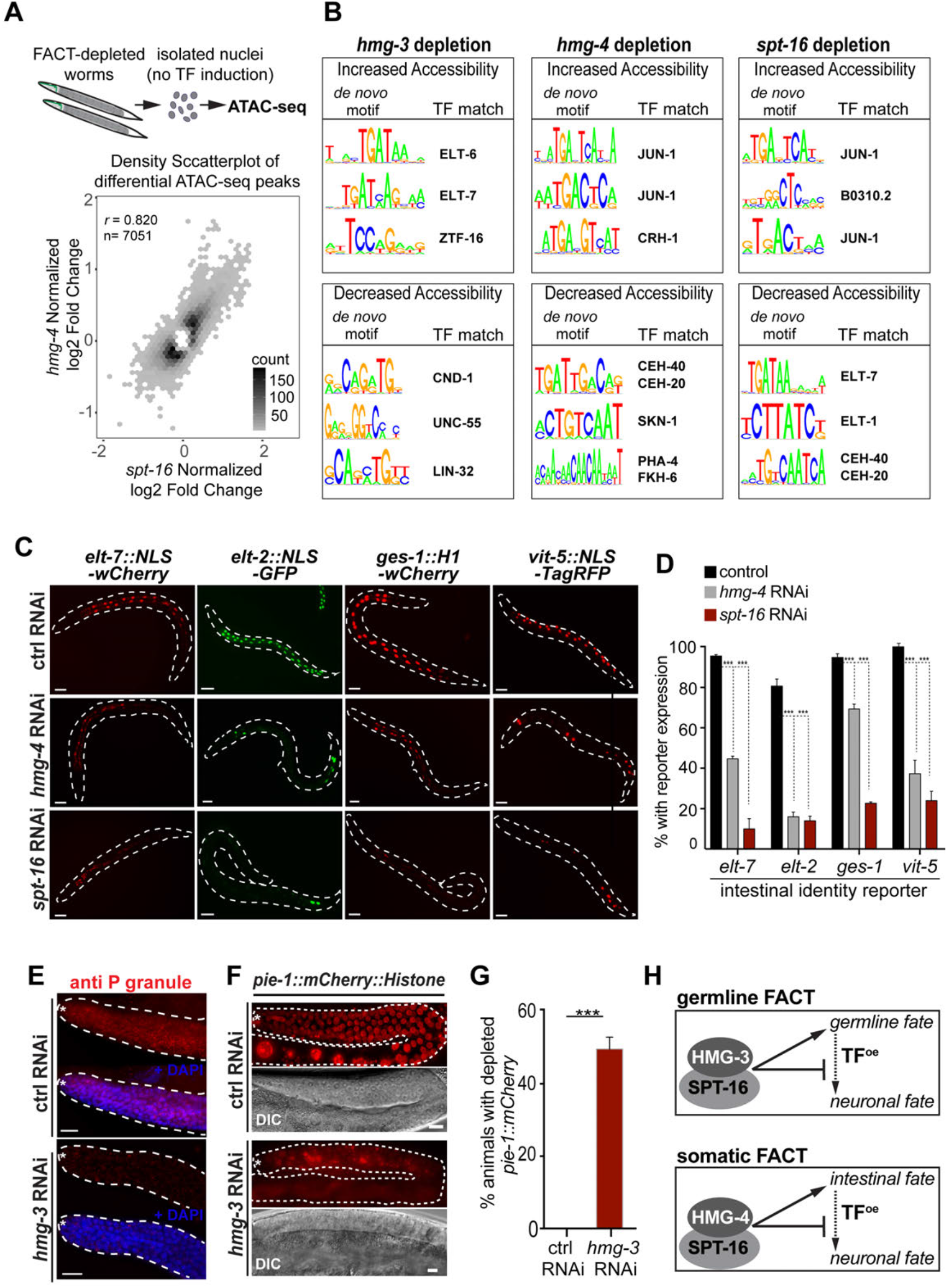
FACT-Depletion Causes Changes in Chromatin Accessibility and Affects Maintenance of Intestinal and Germline Fate. (A) Schematic illustration of ATAC-seq using tsolated nuclei of animals treated with RNAi against FACT subunits without induction. Hexbin density scatterplot of *hmg-4^RNAt^* normalized log2 fold-changes plotted against *spt-16^RNAi^* normalized log2 fold-changes for ATAC-seq peaks that were significantly differential (q < 0.01) in at least one of the conditions. (B) *De novo* motif generation from closing and opening regions in response to *hmg-3^RNAi^, hmg-4^mAi^, and spt-16^RNAi^*. Top 3 enriched motifs with significant matches (details described in Methods) are listed. The full list and orientation of the generated motif relative to the best TF match in the database is provided in Figure S3. (C) RNAi against *hmg-4* or *spt-16* decreases expression of intestinal fate reporters *elt-7::wCherry, elt-2::GFP, ges-1::wCherry* or *vit-5::NLS::tagRFP* Scale bars, 20 μm. (D) Quantification of intestinal fate reporter expression from (C) in *hmg-4^RNAi^* and *spt-16^RNAi^* animals compared to control RNAi. 2-way ANOVA test was used for statistical comparison, *** p<0.001. At least 200 animals were counted for each condition. Error bars represent SEM. (E) Immunostaining of germline-specific P granule in control and *hmg-3^RNAi^* animals. Dashed lines outline the germline, white asterisk indicates the distal tip end of the gonad. Scale bars, 5 μm. (F) The germ cell fate marker *pie-1::mCherry::his-58* is lost in reprogrammed germ cells upon RNAi against *hmg-3*. Dashed lines outline the germline, white asterisk indicates the distal tip end of the gonad. Scale bars, 5 mm. Scale bars, 5 μm. (G) Quantification of *pie-1::mCherry::his-58* loss upon RNAi against *hmg-3* as described in F. Error bars represent SEM. Paired-end student t test was used for statistical comparison, *** p<0.001. (H) Summarizing model: Germline-specific FACT maintains the germ cell fate and block reprogramming to neurons while somatic FACT maintains intestinal gene expression and prevents induction of neuronal genes in the intestine.

ATAC-seq upon RNAi against *hmg-4* and *spt-16* were performed in *glp-4(bn2)* mutant animals that lack a germline when grown at the non-permissive temperature (Beanan and Strome, 1992) to measure chromatin accessibility changes only from somatic cells. A Pearson correlation of 0,820 of log2 fold-changes between sites significantly (q<0.01) changed upon knockdown of *hmg-4^mAl^* or *spt-16^KNAl^* suggests that these factors act together and confirms specificity of the effects (Figure 5A).

Overall, FACT depletion did not lead to a drastic alteration of chromatin accessibility as illustrated by unchanged positional patterns of accessibility around annotated transcription start sites (TSS) and candidate enhancers (Figure S3A and S3B). Nevertheless, *hmg-3^KNAl^* led to 5284 sites becoming more closed and 4747 sites becoming more open, while *hmg-4^m/A^* and *spt-16^KNAl^* caused 1845 and 2472 closing sites as well as 2770 and 2768 sites with increased chromatin accessibility, respectively (Figure 5A and Table S2). While the increase in genomic sites that become less accessible upon FACT depletion is in agreement with the complex’s known function as an activator of gene expression (Hammond et al., 2017; Orphanides et al., 1998), the detection of many increased accessible sites is unexpected and indicates that FACT is also required to prevent ectopic gene expression. This notion is further supported by transcriptome analysis (RNA-seq) of FACT-depleted worms (Figure S4). Depletion of *hmg-3, hmg-4* or *spt-16*, leads to down regulation but also up-regulation of several genes-e.g. 1679 down-regulated and 1948 up-regulated genes upon *spt-16* RNAi (Figure S4).

In order to gain insight whether chromatin sites that change accessibility upon FACT depletion are enriched for binding of TF families we performed *de novo* motif analysis in ATAC-seq peaks. Notably, closing chromatin sites in *hmg-3^mAl^* animals are enriched for bHLH family motifs matching that of the NeuroD-related TF CND-1, while motifs for GATA family TFs matching those for ELT-6 and ELT-7 are enriched at opening chromatin sites (Figures 5B and S3C). Considering that germ cells can be reprogrammed to neurons upon *hmg-3* RNAi it is surprising that potential binding sites for a neuroblast-specifying TF such as CND-1(Hobert, 2010) become less accessible. This could indicate that during conversion to neurons germ cells bypass early developmental routes of neuron specification. In *spt-16^mAl^* animals motifs matching the master regulator GATA TF of intestinal fate ELT-7 (Sommermann et al., 2010) is enriched at closing sites. In *hmg-4^mAl^* animals, motifs for the bZIP TF family matching SKN-1 and FoxA TF family matching PHA-4, which co-regulate the intestinal fate (Azzaria et al., 1996; Horner et al., 1998; McGhee, 2007), are enriched in closing sites (Figures 5B and S3C). Decreased accessibility for TFs that regulate intestinal gene expression might reflect a loss of intestinal cell fate maintenance upon RNAi against *hmg-4* or *spt-16*. Therefore, we performed postembryonic RNAi against *hmg-4* or *spt-16* and examined the expression of intestinal genes using four different intestine-specific reporters as well as immunostaining for the intestine-specific proteins ELT-2 (GATA TF) and IFB-2 (intermediate filament protein) (Figure S5). Somatic FACT depletion caused a loss of intestinal gene expression (Figures 5C–5D and S5) suggesting that FACT is required to maintain the gut cell identity. Similarly, *hmg-3* RNAi caused decreased levels of germline P-granules and significant reduction of the germline-specific *pie-1* reporter expression (Figures 5E – 5G) indicating that FACT also maintains the germline fate. In summary (Figure 5H), our results suggest that loss of germline or soma-specific FACT leads to both-decreased as well as increased chromatin accessibility for TFs and impaired maintenance of germline and intestinal cell fate identities, respectively.

### FACT is a Reprogramming Barrier in Human Fibroblasts

As a chromatin regulator, FACT is highly conserved through evolution, suggesting that its role in cellular reprogramming may also be conserved. Therefore, we tested whether FACT depletion in human fibroblasts enhances reprogramming using a cell line (hiF-T) that shows reprogramming to iPSCs only with very low efficiency (Cacchiarelli et al., 2015). The hiF-T fibroblasts allow doxycycline (DOX)-inducible expression of the Yamanaka TFs OCT4, SOX2, KLF4, and C-MYC (OSKM) derived from a stable transgene that ensures expression of comparable OSKM levels in repeat experiments (Cacchiarelli et al., 2015) (Figure 6A). The human FACT subunits SSRP1 and SUPT16H were efficiently depleted using transfection of short interfering RNAs (siRNAs) for up to 4 days while transcript levels started recovering after 7 days (Figure S6A). OSKM induction (+DOX) 48h after siRNA transfection against SUPTH16 and SSRP1 considerably increased the numbers of iPSC colonies – approximately by 90% upon SUPTH16 depletion (Figures 6B and 6C). FACT depletion does not de-repress the DOX-inducible OSKM transgenes (Figure S6B) excluding the possibility of reprogramming enhancement simply due to affecting the OSKM cassette. These colonies show strong expression of several pluripotency markers including NANOG, SSEA4 and Tra-1-60 (Park et al., 2008) (Figure S6C and S6D). Additionally, pluripotency was confirmed in a physiological context by transplanting the derived iPSCs into mice which then formed teratomas containing tissues from all three germ layers (Brivanlou et al., 2003; Hentze et al., 2009; Kooreman and Wu, 2010) (Figure 6D). Next, we asked whether FACT depletion in human fibroblast also enhances lineage reprogramming to neurons (Figure 6E).

**Figure 6.**
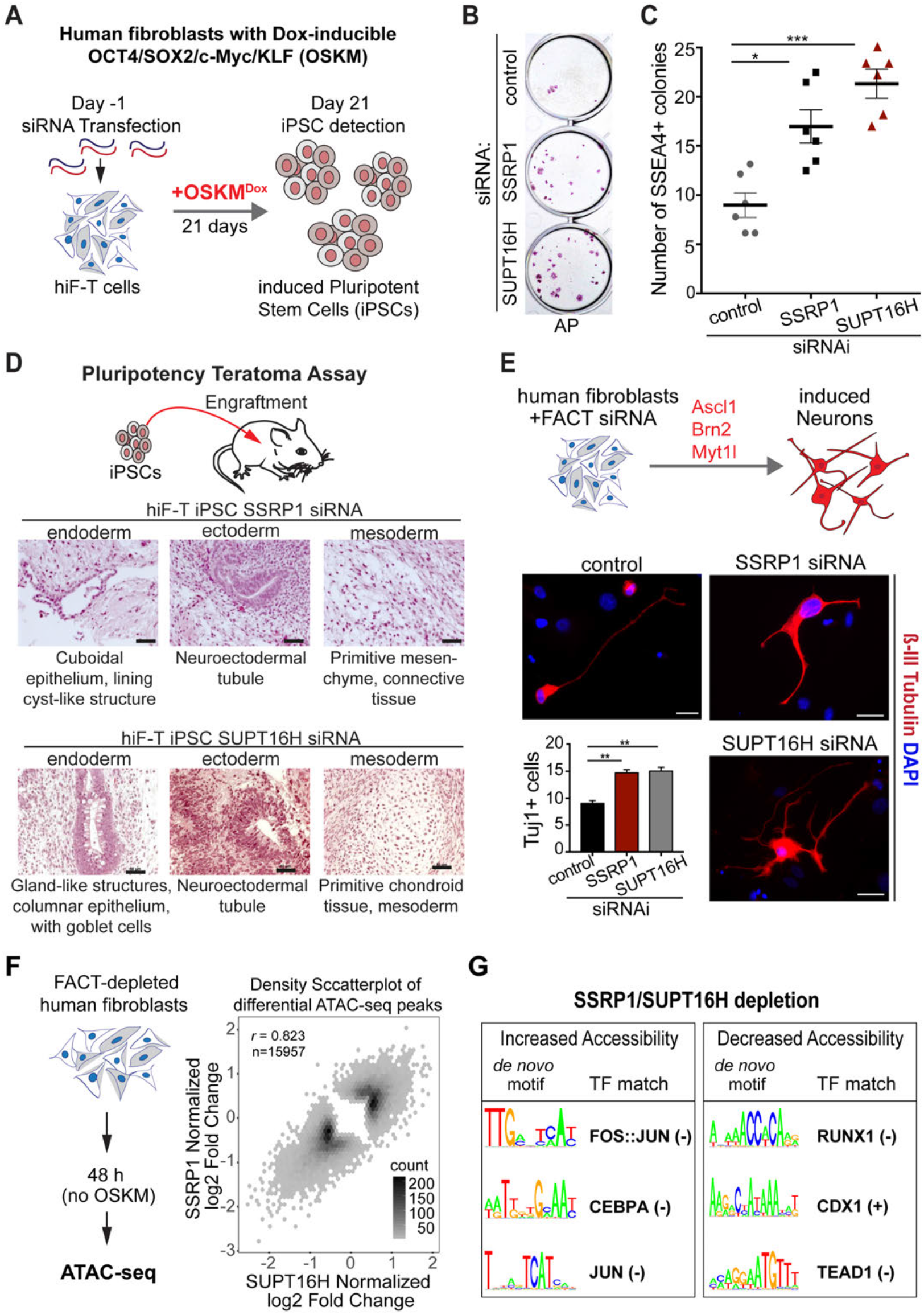
FACT Depletion Increases Efficiency of Reprogramming Human Fibroblasts. (A) Schematic showing procedure with human secondary fibroblasts carrying a doxycycline (DOX)-inducible, poly-cistronic human OCT4/SOX2/KLF4/c-MYC (OSKM) cassette (Cacchiarelli et al., 2015) that were transfected with siRNAs against human FACT subunit encoding genes SSRP1 and SUPTH16 before DOX induction. (B) Alkaline phosphatase (AP) staining of stem cell colonies 21 days after DOX treatment. Control experiment is scrambled siRNA (see Methods). (C) Quantification of iPSC colonies from 6 replications based on SSEA4 immunostaining as shown in Figure S6 for each knock-down condition. Paired student t test was used for statistical comparison, *p<0.05, **p<0.01, *** p<0.001. (D) Histological characterization of teratomas derived from grafting SSRP1 or SUPT16H depletion-derived iPSCs in mice. Teratomas reached 1.5 cm^2^ in size on average after 51 days (SSRP^RNAi^ iPSCs) or 70 days (SUPT16H^RNAi^) and show tissues of all three germ layers (ectoderm, endoderm, mesoderm). Scale bars, 50 μm (E) Forced expression of the TFs Ascl1, Brn2, Mytl1 (BAM) in human fibroblasts (NHDF cells) and SSRP1 or SUPT16H depletion enhances reprogramming of fibroblasts to neurons by approximately 50%. Beta-III tubulin-positive neurons derived from FACT-depleted fibroblasts showed a higher degree of projection complexity compared to control cells. One-way ANOVA test was used for statistical comparison, ** p<0.01. (F) ATAC-seq was performed in hiF-T cells without OSKM induction 48h after FACT depletion. Hexbin density scatterplot of SSRP1^RNAi^ normalized log2 fold-changes plotted against SUPT16H^RNAi^ normalized log2 fold-changes for those ATAC-seq peaks that were significantly differential (q < 0.01) in at least one of the two conditions. (G) *De novo* motif generation (see Material and Methods) in opening and closing regions upon SSRP1^RNAi^ and SUPT 16H^RNAi^. Top 3 enriched motifs for each indicated set are given.

Although transduction efficiency was relatively low (15%) our results suggest that forced expression of the TFs Ascl1, Brn2, Mytl1 that were shown previously to induce conversion of fibroblasts into neurons (Vierbuchen et al., 2010) was able to increase the neuronal conversion up to 50% upon FACT depletion as compared to the control (Figure 6E). Interestingly, the beta-III tubulin-positive neurons derived from FACT depleted fibroblasts show a higher degree of projection complexity than controls cells (Figure 6E). This could indicate that the reprogramed cells are generated earlier or mature faster upon FACT depletion. Taken together, FACT depletion in human fibroblasts enhances their reprogramming to iPSCs and neurons demonstrating that FACT’s role as a safeguard of cellular identities is conserved in human cells.

### FACT Depletion Has Analogous Effects on Chromatin in Humans and *C. elegans*

In order to examine whether FACT depletion causes similar changes in chromatin accessibility as seen in *C. elegans* we performed ATAC-seq using FACT-depleted fibroblasts (Figure 6G). A Pearson correlation of 0.823 of log2 fold-changes between sites significantly (q < 0.01) changed upon knockdown of SSRP1 or SUPTH16 confirms that both factors act together (Figure 6F). As in *C. elegans*, FACT depletion did not lead to a drastic alteration of chromatin accessibility as illustrated by unchanged positional patterns of accessibility around annotated transcription start sites (TSS) and annotated enhancers (Figure S6E and S6F). Nevertheless, in fibroblasts 4467 genomic sites commonly changed their chromatin accessibility upon treatment with either SSRP1 or SUPT16H siRNA: 2846 sites became more closed while 1621 genomic sites gained higher accessibility (Figure 6F and Table S3). As in *C. elegans*, the increase in chromatin accessibility upon FACT depletion is unexpected due to FACT’s established function as a positive regulator of gene expression (Orphanides et al., 1998). A striking similarity upon FACT depletion in human cells as well in *C. elegans* is the enrichment for motifs of the conserved bZIP TF JUN/JUN-1 at chromatin sites with increased accessibility (Figures 6H and S6F). As part of the AP-1 TF complex in humans c-Jun is involved in numerous cellular processes including cell growth and proliferation (Shaulian, 2010) by, for instance, inhibiting p53 (Eferl et al., 2003; Schreiber et al., 1999). In *C. elegans* the transcripts of *jun-1* are enriched in the intestine (McGhee et al., 2007) but its role in this tissue is unclear. However, a recent study suggests that JUN-1 might cooperate with GATA TFs such as ELT-2 during regulation of the innate immune response in the intestine (Yang et al., 2016). Furthermore, FACT depletion leads to decreased chromatin accessibility for CDX TFs (Figure 6G) which are important for maintaining and inducing intestinal gene expression (Fujii et al., 2012; San Roman et al., 2015), an effect that is similar to loss of somatic FACT in *C. elegans* (Figures 5B – D). In fibroblasts, the most enriched motif at closing chromatin sites matches that for the Runt-related transcription factor 1 (RUNX1) (Figures 6G and S6F). Interestingly, RUNX1 depletion has been shown to enhance reprogramming (Chronis et al., 2017) suggesting that decreased chromatin accessibility for RUNX1 binding sites upon FACT depletion could have analogous effects. Likewise, decreased accessibility for sequences matching the TEAD family TF motif) (Figures 6G and S6F) could affect the TEAD/HIPPO pathway – another previously reported reprogramming barrier (Qin et al., 2012). Furthermore, increased accessibility for the Cebp family TFs (Figures 6H and S6F) might trigger their redistribution which has been shown to enhance reprogramming in the context of mouse embryonic fibroblasts (Chronis et al., 2017; Di Stefano et al., 2016). Overall, FACT’s combined role in maintaining gene expression and preventing chromatin accessibility for some TFs such as Jun/JUN-1 in human fibroblasts and *C. elegans* seems conserved and might be essential to prevent deviant expression of genes belonging to other cell identities in both species. This way FACT safeguards cell identities and antagonizes reprogramming by TFs in *C. elegans* and Human.

## Discussion

The identification of the highly conserved histone chaperone FACT as a barrier for cellular conversion supports a recent notion by Alvarado and Yamanaka (Alvarado and Yamanaka, 2014) that cell fate maintenance factors have not been widely explored but might provide new avenues for reprogramming. We identified FACT as an impediment for cellular reprogramming in an unbiased genetic screen using *C. elegans*, which has two SSRP1 orthologs, HMG-3 and HMG-4, giving rise to germline and soma-specific FACT isoforms. Such tissue-specific isoforms have so far not been reported in any other species but organisms such as Zebrafish also contain two different SSRP1 homologs (treefam.org) raising the possibility that alternative FACT isoforms might safeguard different tissues in other species as well.

Overall, the discovery of FACT as a cellular reprogramming barrier was highly unexpected since FACT is predominantly known as a positive regulator of gene expression by facilitating transcription (Orphanides et al., 1998) (reviewed by (Hammond et al., 2017; Reddy et al., 2017)). It is conceivable that loss of FACT has two consequences: reduced gene expression required to maintain cell identities, such as the intestinal and germ cell fate in *C. elegans*, and increased accessibility at chromatin sites that are otherwise repressed. Strikingly, motifs for the AP-1 TF Jun/JUN-1 are enriched in both human fibroblasts and *C. elegans* at opening chromatin sites upon FACT depletion indicating a remarkable conservation of gene regulation networks. Though it remains unclear whether increased permissiveness for JUN-1 activity could promote changes in cell identities in *C. elegans*, in mammals c-Jun is an inhibitor of p53 during development and in tumors (Eferl et al., 2003; Schreiber et al., 1999) indicating a possible mechanism for reprogramming enhancement (Hanna et al., 2009). However, a recent study showed that in mouse embryonic fibroblasts (MEFs) c-Jun impedes reprogramming to iPSCs (Liu et al., 2015) suggesting that c-Jun has either dichotomous roles for regulating cell identities as previously shown for other reprogramming regulators (Rais et al., 2013), or it has context dependent activities for regulating reprogramming in MEFs versus human cells. Generally, the observed dynamics of accessible sequence content upon FACT depletion could poise human fibroblasts for reprogramming due to a preponderance of regulatory effects that favor reprogramming such as increased accessibility for CEBP TF binding sites s together with decreased binding of RUNX1 and TEAD/HIPPO TFs. How FACT depletion leads to chromatin sites that become more accessible remains to be determined. Since FACT is also required for re-establishing the nucleosome signature after RNA polymerase II passage (Belotserkovskaya et al., 2003; Jamai et al., 2009) it is possible that lack of FACT leads to more accessible chromatin due to persistence of nucleosome-depleted sites. More indirectly, decreased gene expression upon loss of FACT could obliterate the insulation of active and repressed chromatin regions which has been previously implied for cell identity genes (Dowen et al., 2014). Nevertheless, FACT’s activity as a maintenance factor of gene expression and chromatin accessibility suggests that FACT is a genuine safeguarding factor of cellular identities.

For application aspects, transient FACT depletion in human cells is sufficient to enhance reprogramming, thereby providing new avenues for clinical approaches. It has been shown that short-term OSKM expression is sufficient for *in vivo* reprogramming in mice thereby preventing formation of tumors (Ocampo et al., 2016). In general, the combination of transient depletion of reprogramming barriers and short-term forced-expression of TFs will have less deleterious effects while ensuring efficient reprogramming.

Seen in a broader context, FACT being an impediment for converting cellular identities in *C. elegans* and human cells exemplifies that reprogramming barriers are evolutionarily conserved, a phenomenon that is also reflected by the previously identified barrier for germ cell reprogramming LIN-53 in *C. elegans* – CAF-1p48/RBBP7 in mammals (Tursun et al., 2011). The LIN-53-containing histone chaperone CAF-1 acts as a barrier during reprogramming of mouse fibroblasts (Cheloufi et al., 2015) indicating that the underlying mechanisms are deeply conserved. Overall, our study demonstrates the versatility of *C. elegans* as a gene discovery model for identifying novel and unanticipated reprogramming barriers. Since reprogrammed cells have gained importance for tissue replacement therapies, exploring new avenues that reveal unexpected roadblocks to cell fate reprogramming may facilitate the generation of human tissues for therapeutic applications.

## Author Contributions

Conceptualization, B.T. and E.K.; Methodology, B.T., E.K., S.A.L., and S.D.; Investigation, E.K., A.O., G.B, S.S., A.S., M.H., B.T., B.U., A.A., and S.D.; Validation, B. T., E.K., S.A.L., and S.D.; Writing – Original Draft, B.T.; Writing – Review and Editing, B.T., E.K., S.A.L., B.U., A.A., and S.D.; Funding Acquisition, B.T.; Resources, B. T., S.A.L., and S.D., Visualization, B.T., E.K., S.A.L., and S.D.; Supervision, B.T., S.A.L., and S.D.; Project Administration, B.T.; Funding Acquisition, B.T.

## Acknowledgments

We thank Oliver Hobert, Roger Pocock, Hannes Bülow, Luisa Cochella and Claude Desplan for critical reading of the manuscript and comments. We thank Sergej Herzog, Alina El-Khalili, Norman Krüger and Antje Hirsekorn for technical assistance. Also, we thank Uwe Ohler and Neelanjan Mukherjee for useful discussion and support with respect to Bioinformatics. We thank members of the Tursun group for comments on the manuscript and Dr. Mikkelsen for providing hiF-T cells, CGC for worm strains (supported by the NIH). This work was sponsored by the ERC-StG-2014-637530 and ERC CIG PCIG12-GA-2012-333922 and is supported by the Max Delbrueck Center for Molecular Medicine in the Helmholtz Association.

---

**Figure S1.**
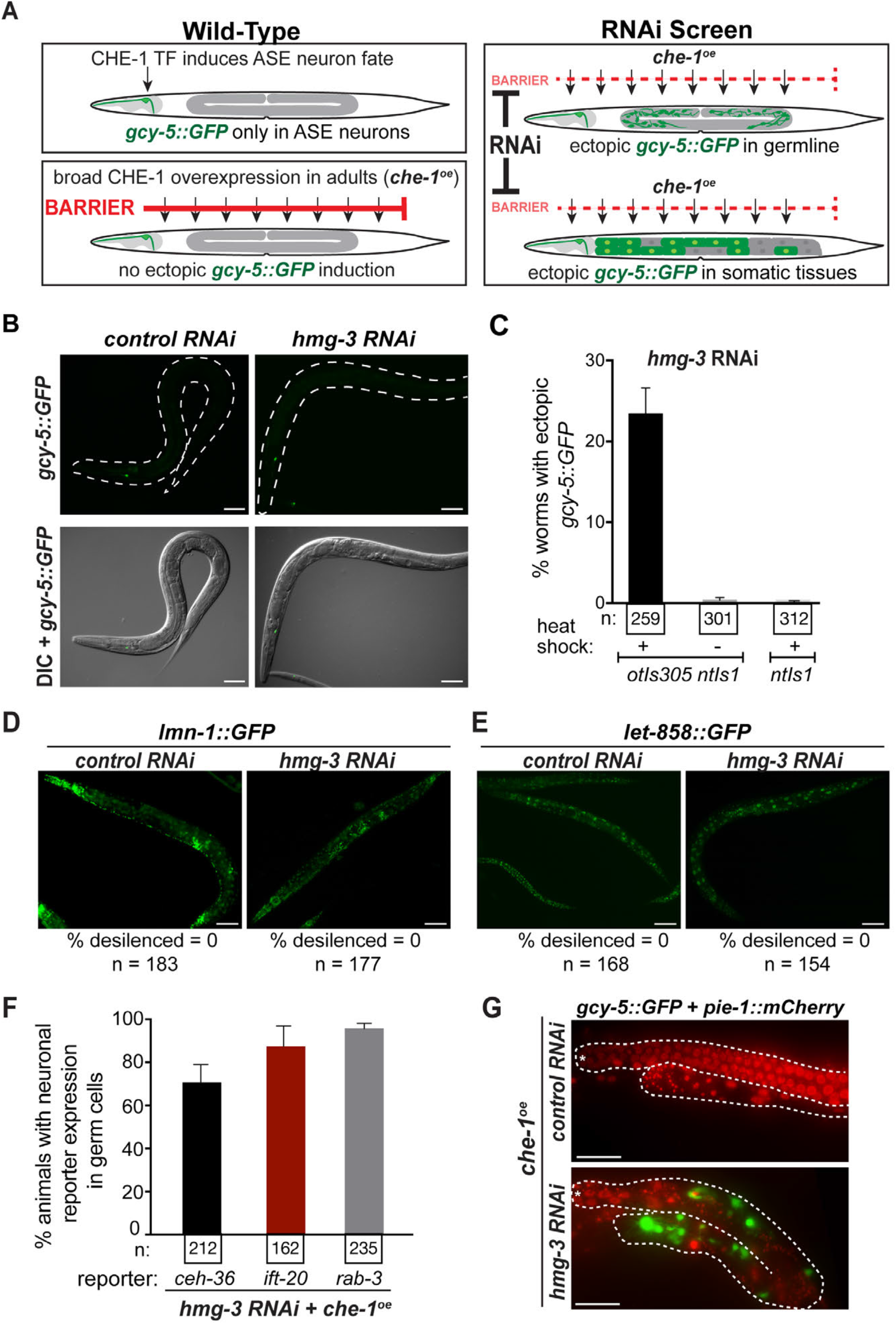
Assessment of Ectopic *gcy-5::GFP* Induction in Germ Cells Upon RNAi Against *hmg-3*, Related to Figure 1. (A) Schematic representation of transgenic animals that express the *gcy-5::GFP* reporter for ASE neuron fate induced by zinc-finger TF CHE-1. Ubiquitous mis-expression of CHE-1 *(che-1^oe^)* in adults does not induce ectopic *gcy-5::gfp*. An RNAi screen to identify factors that prevent ectopic *gcy-5::GFP* by *che-1^oe^* was carried out. (B) Representative images of *hmg-3^KNAl^* animals with and without *che-1^oe^* (no *hs::che-1* construct in the background), or with only *gcy-5::GFP (ntIs1)* construct in the background. *gcy-5::GFP* is not induced upon *hmg-3* RNAi without *che-1^oe^*. Scale bars, 20 μm. (C) Quantification of *gcy-5::GFP* induction in *hmg-3^mAl^* with and without *che-1^oe^* (no *hs::che-1* construct in the background) or with only *gcy-5::GFP (ntIs1)* construct in the background. Error bars represent SEM. (D and E) Representative images of *lmn-1::GFP* (D) and *let-858::GFP* (E) animals after *hmg-3* RNAi. Neither expression of *lmn-1::GFP* nor expression of *let-858::GFP* changes upon FACT depletion. Scale bars, 20 μm. (F) Quantification of neuronal markers *ceh-36::RFP, ift-20::NLS::RFP* and *rab-3::NLS::RFP* in *gcy-5::GFP* positive germline of *hmg-3^KNAl^* animals. Error bars represent SEM. (G) The germ cell fate marker *pie-1::mCherry::his-58* is lost in reprogrammed germ cells. Dashed lines indicated the outline of the gonad, Asterisk labels distal tip of the germline. Scale bars, 5 μm.

**Figure S2.**
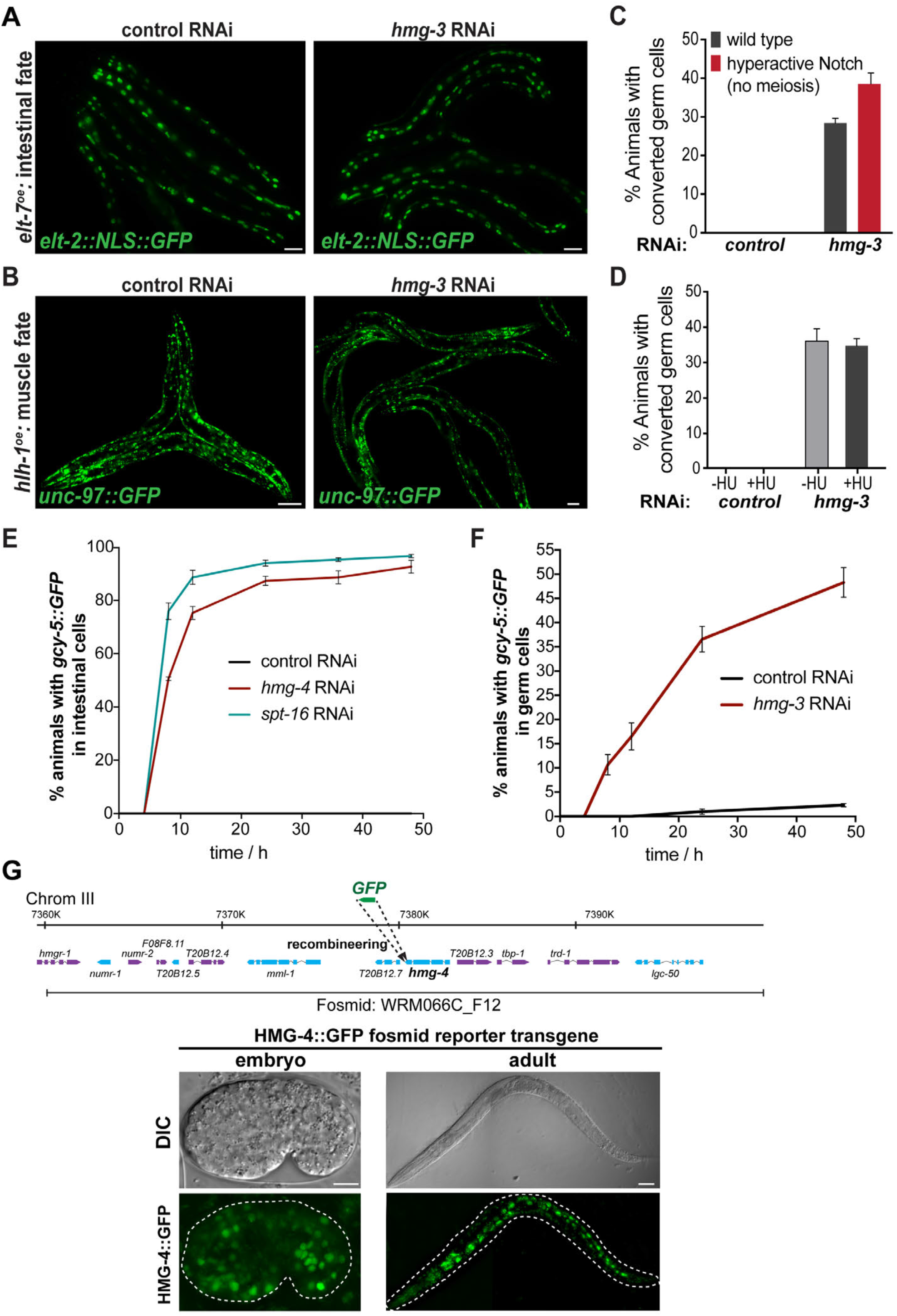
Depletion of HMG-3 Allows GABA and ASE Neuron-Fate, but not Muscle or Intestinal Fate Induction in Germ Cells, Related to Figures 2 and 3. (A and B) Representative pictures showing that overexpression of (A) intestinal or (B) muscle fate-inducing TFs do not convert germ cells into gut or muscle cells in *hmg-3^KNAl^* animals. Scale bars, 20 μm. (C) Germ cell conversion is not lost in the *glp-1(gf)* mutant background which lacks meiotic germ cells, but retains mitotic germ cells. More than 150 animals were counted for each condition. Error bars represent SEM. (D) Reprogramming is independent of cell cycle. Phenotype penetrance remained unchanged after HU-mediated cell cycle arrest (-HU, no treatment, +HU, 6 hr HU treatment). More than 150 animals were counted. Error bars represent SEM. (E) Time course experiment *gcy-5::GFP* induction in *hmg-4^mAl^* and *spt-16^KNAl^* animals. Time in h after induction of *che-1^oe^*. More than 150 animals were counted for each time point. Error bars represent SEM. (F) Time course experiment of *gcy-5::GFP* induction in *hmg-3^KNAl^* animals. Time in h after induction of *che-1^oe^*. More than 150 animals were counted for each time point. (G) Schematic of recombineered fosmid containing the *hmg-4* locus with GFP and representative images of *hmg-4::GFP^fosĦld^* expression in young adult and comma stage embryo. DIC = Differential interference contrast. Scale bars, 20 μm.

**Figure S3.**
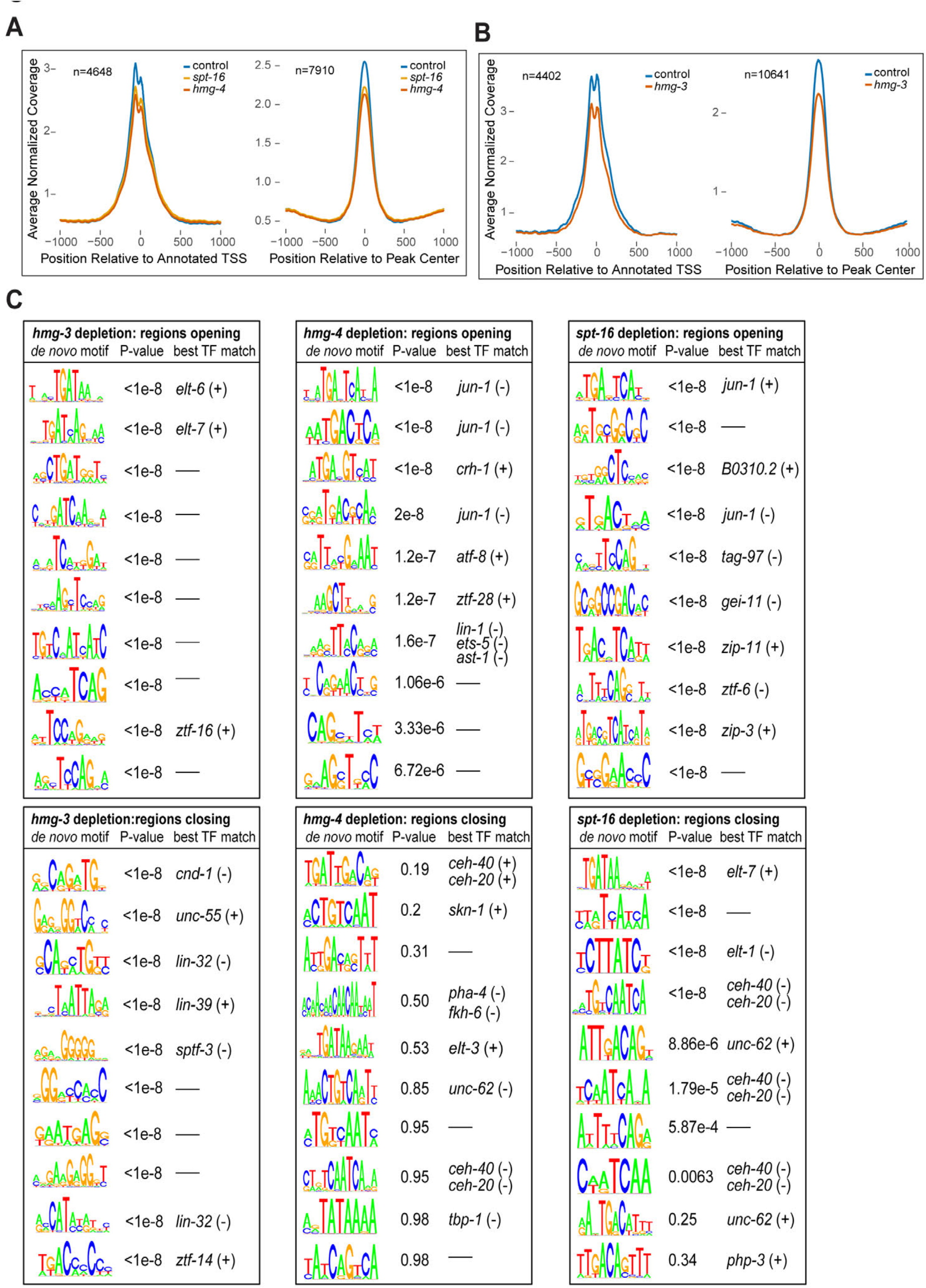
ATAC-seq Analysis of Worms After RNAi Against *hmg-3, hmg-4* and *spt-16*, Related to Figure 5. (A and B) Distribution of normalized ATAC-seq coverage relative to gencode annotated transcription start sites (TSS) that intersected ATAC-seq peaks and relative to midpoints of candidate enhancers in (A) hmg-3^RNAi^ or (B) hmg-4^RNAl^ and *spt-16^RNAl^*. (C) *De novo* motif generation (see Material and Methods) in opening or closing regions upon *hmg-3^KNAl^, hmg-4^RNAl^* and *spt-16^mAl^*. Top 10 enriched motifs for each indicated set together with the p value and the best TF match are given. The orientation of the generated motif relative to the best TF match in the database is indicated in parentheses.

**Figure S4.**
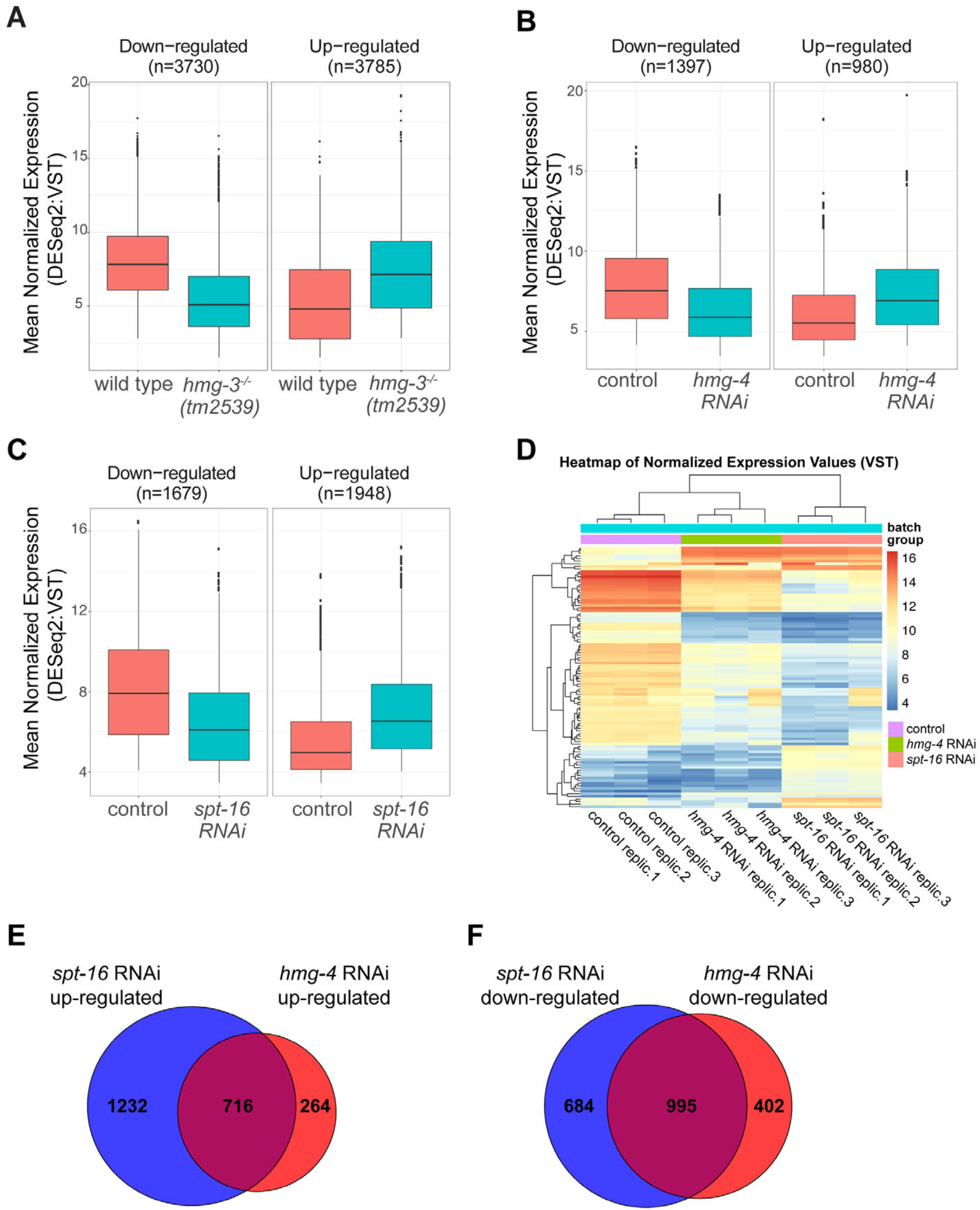
RNA-seq Analysis of Worms After RNAi Against *hmg-3, hmg-4* and *spt-16*, Related to Figure 5. (A – C) Expression changes in *C. elegans* in hmg-3-depleted background *(hmg-3(tm2539))* (A), and RNAi against *hmg-4* (B) and *spt-16* (C). 3730 genes were down-regulated and 3785 genes were up-regulated in hmg-3-depleted background *(hmg-3(tm2539))* (A). We detected 1397 up-regulated and 980 down-regulated genes after RNAi against *hmg-4* (B), and 1679 and 1948 up-regulated and down-regulated genes, respectively in *spt-16^RNAl^* animals. Up/down regulated genes are detected based on the differential expression criteria of adjusted p-value of at least 0.1 and at least two-fold increase or decrease in expression levels in relation to the control samples. Transcription levels of these up-and downregulated genes are represented as boxplots. (D) A heatmap of unsupervised hierarchical clustering of the top 100 genes with most variant gene expression across control, *hmg-4* and *spt-16* RNAi samples shows that *hmg-4^mAt^* and *spt-16^KNAl^* are more similar to each other than the control. Independently generated biological replicates clustered together (individual samples and batches are indicated). (E and F) Venn-diagram showing overlap of (E) up-regulated and (F) down-regulated genes (numbers given) in *hmg-4^RNAl^* and *spt-16^RNAl^* animals.

**Figure S5.**
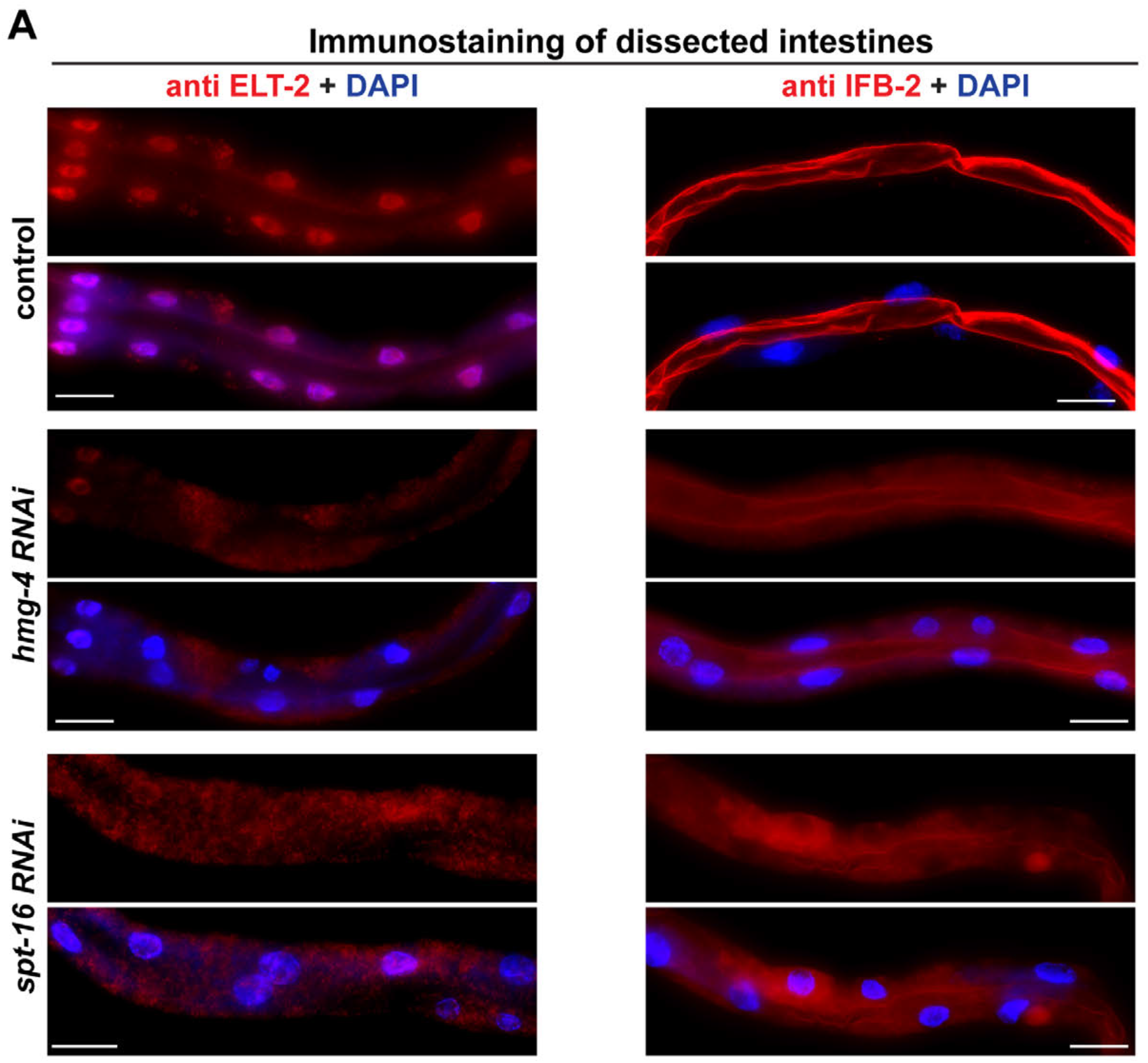
Immunostaining for Intestinal Proteins upon somatic FACT depletion, Related to Figure 5. Intestines from animals treated with RNAi against *hmg-4* and *spt-16* after embryonic development were dissected and immunostained for the gut-specific GATA TF ELT-2 and intestine-specific intermediate filament protein IFB-2. Scale bars, 5 μm.

**Figure S6.**
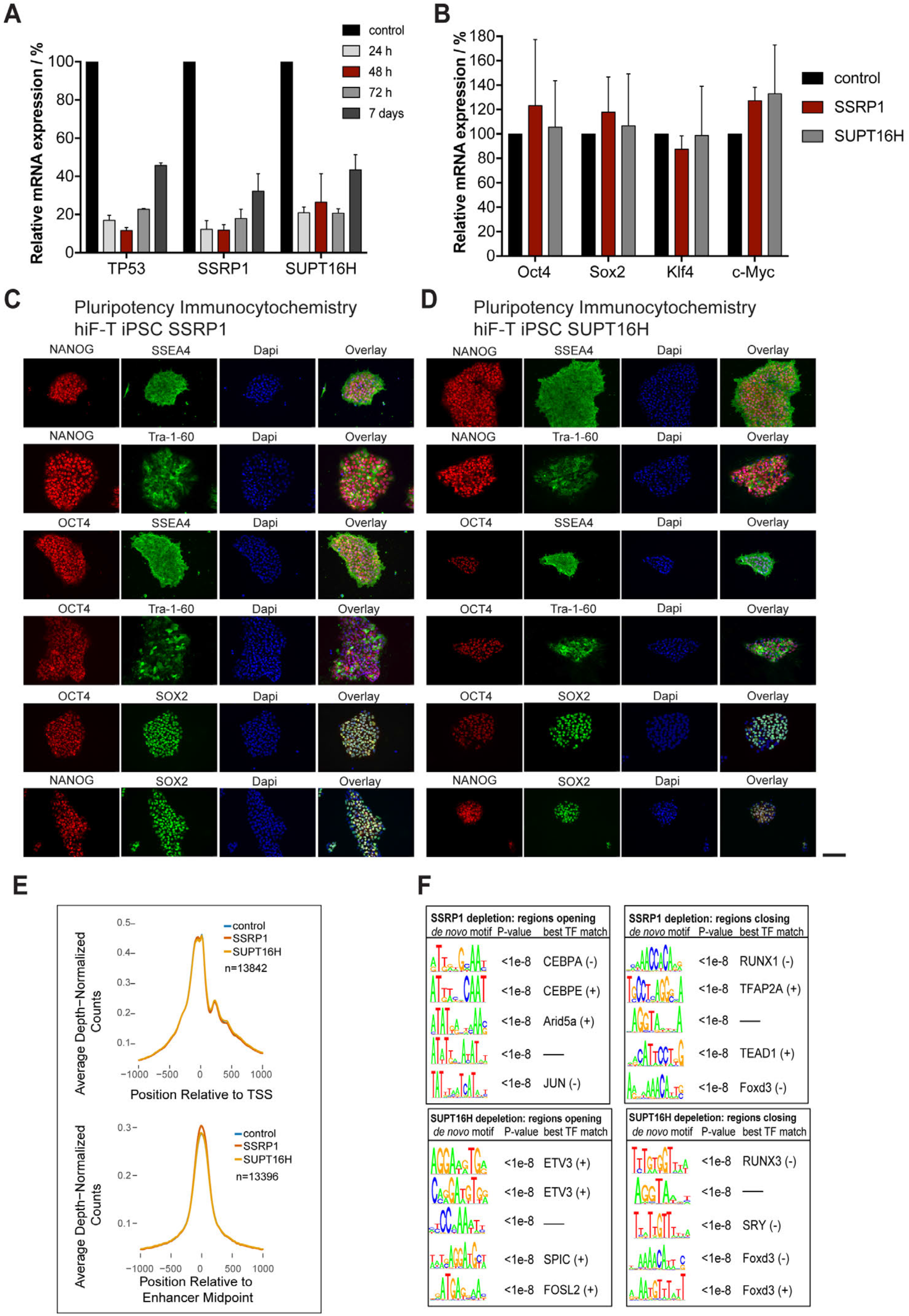
FACT siRNA Knockdown efficiency and Pluripotency Marker Expression in FACT-Depletion Derived iPSCs, Related to Figure 6. (A) Quantitative RT-PCR (qRT-PCR) analysis to confirm knock-down of SSRP1, SUPT16H and TP53 at 24 h, 48 h, 72 h and 7 days after transfection with siRNAs. (B) qRT-PCR analysis for expression of levels of Oct4, Sox2, Klf4, and c-Myc 48 hours after SSRP1 or SUPT16H depletion. Gene expression levels were normalized to GAPDH expression levels and compared to control siRNA. Error bars represent SD. (C and D) Representative images of antibody staining for NANOG, OCT4, SOX2 SSEA-4 and Tra-1-60 in iPSC colonies derived from SSRP1 (C) or SUPT16H (D) depleted hiF-T cells. Scale bars, 25 μm. E) Distribution of normalized ATAC-seq coverage relative to gencode annotated transcription start sites (TSS) that intersected ATAC-seq peaks and relative to midpoints of FANTOM enhancers that intersected ATAC-seq peaks in control, SSRP1^RNAi^, and SUPT16H^rna1^ (F) *De novo* motif generation (see Material and Methods) in closing and opening regions upon SSRP1^RNAi^ (E) and SUPT16H^RNAl^ (F) Top 5 enriched motifs for each indicated set together with the p value and the best TF match are given. The orientation of the generated motif relative to the best TF match in the database is indicated in parentheses.

**Table S1. ATAC-seq basic read statistics, Related to Figures 5, 6 and S5 and S6**

**Table S2. Common Differences in *hmg-4* / *spt-16* and SSRP1 / SUPT16H knockdown Analyzed by ATAC-seq, Related to Figures 5, 6 and S5 and S6**

**Table S3. Differences in *hmg-3* and *hmg-4* knock-down analyzed by ATAC-seq, Related to Figure 6 and S6**

## Methods

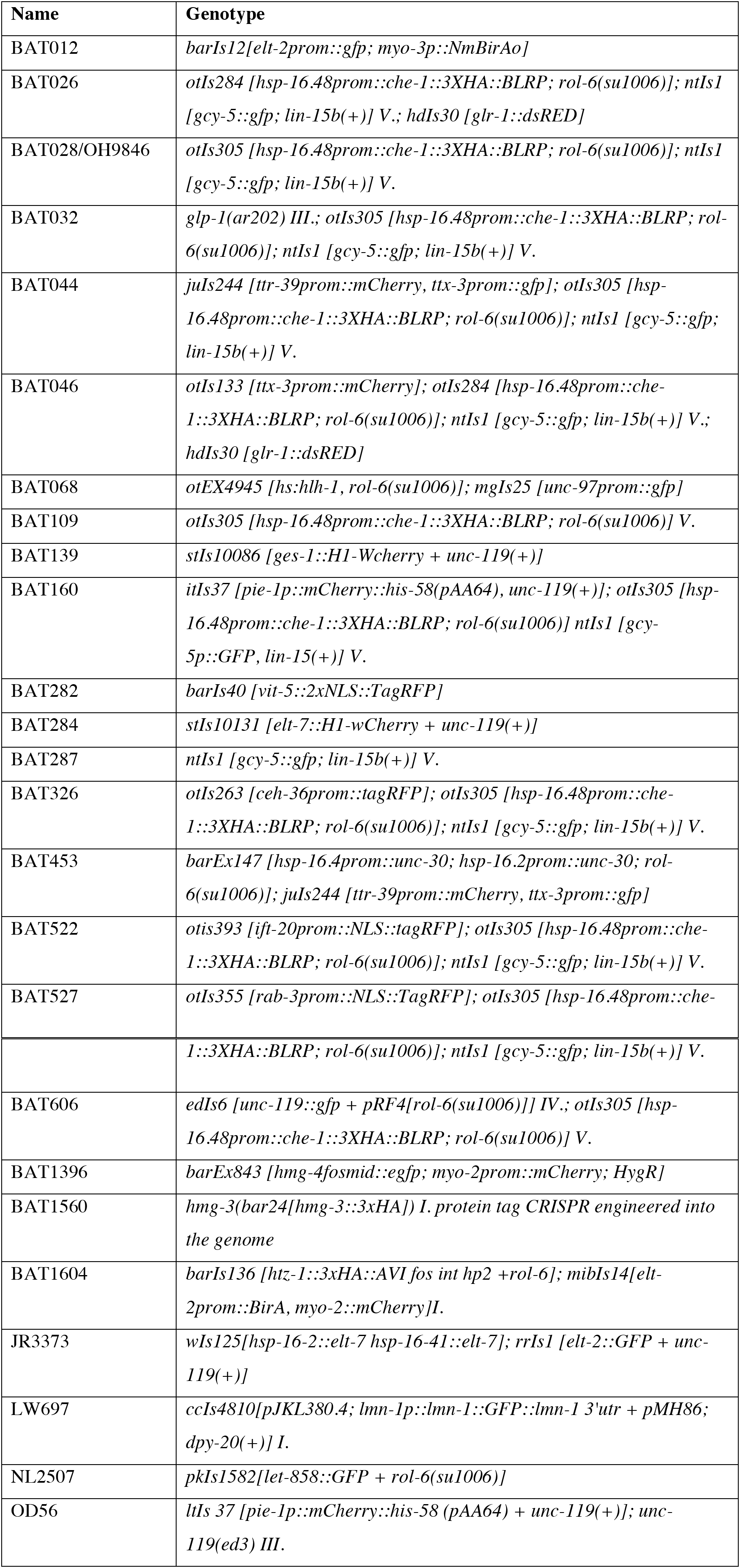
*C. elegans* strains used in the study.

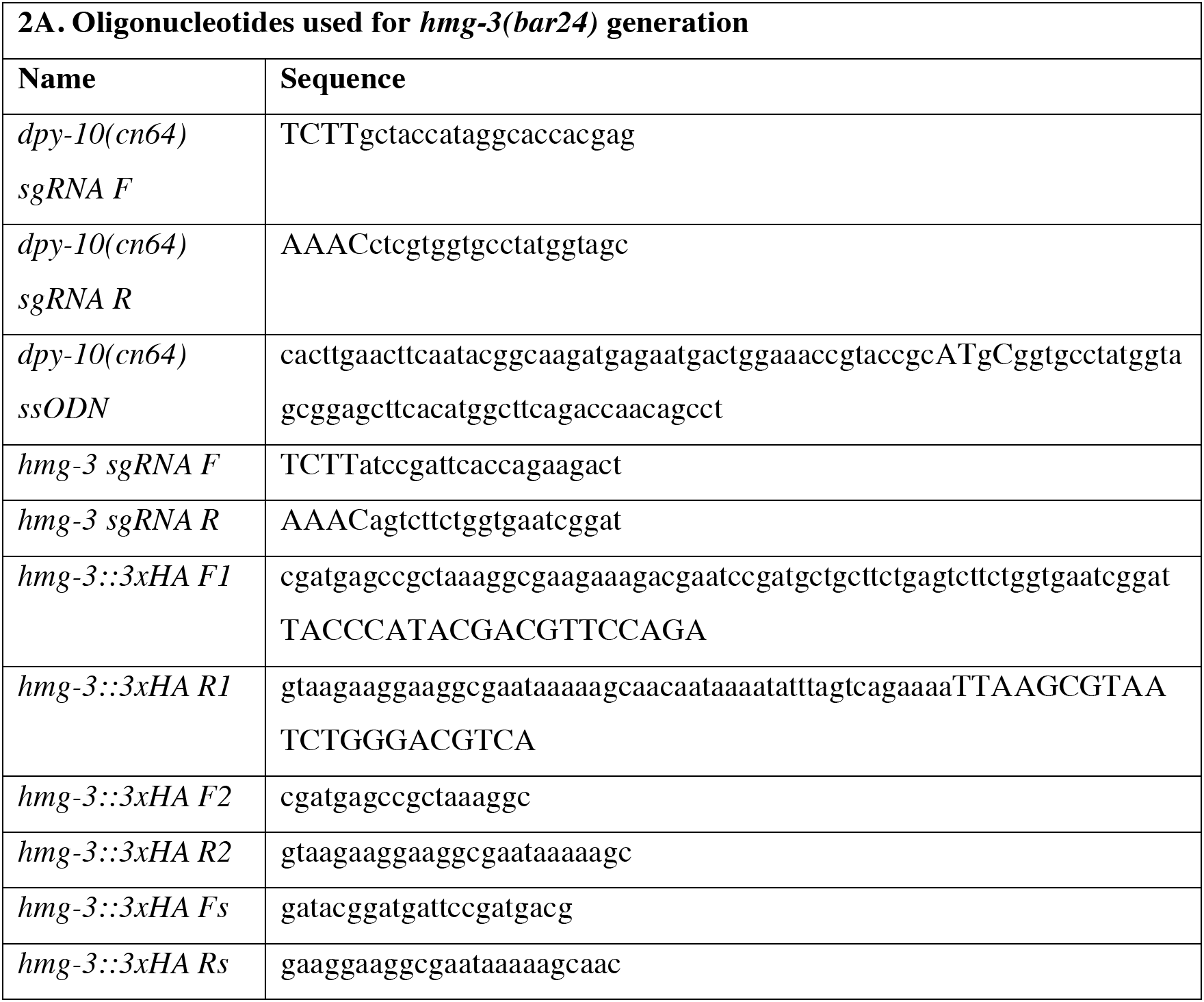
Oligonucleotides used for *hmg-3(bar24)* generation by CRISPR/Cas9 and smFISH probe sequences.

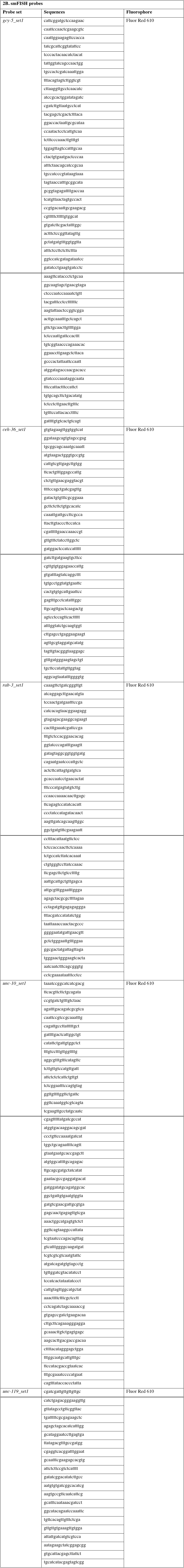

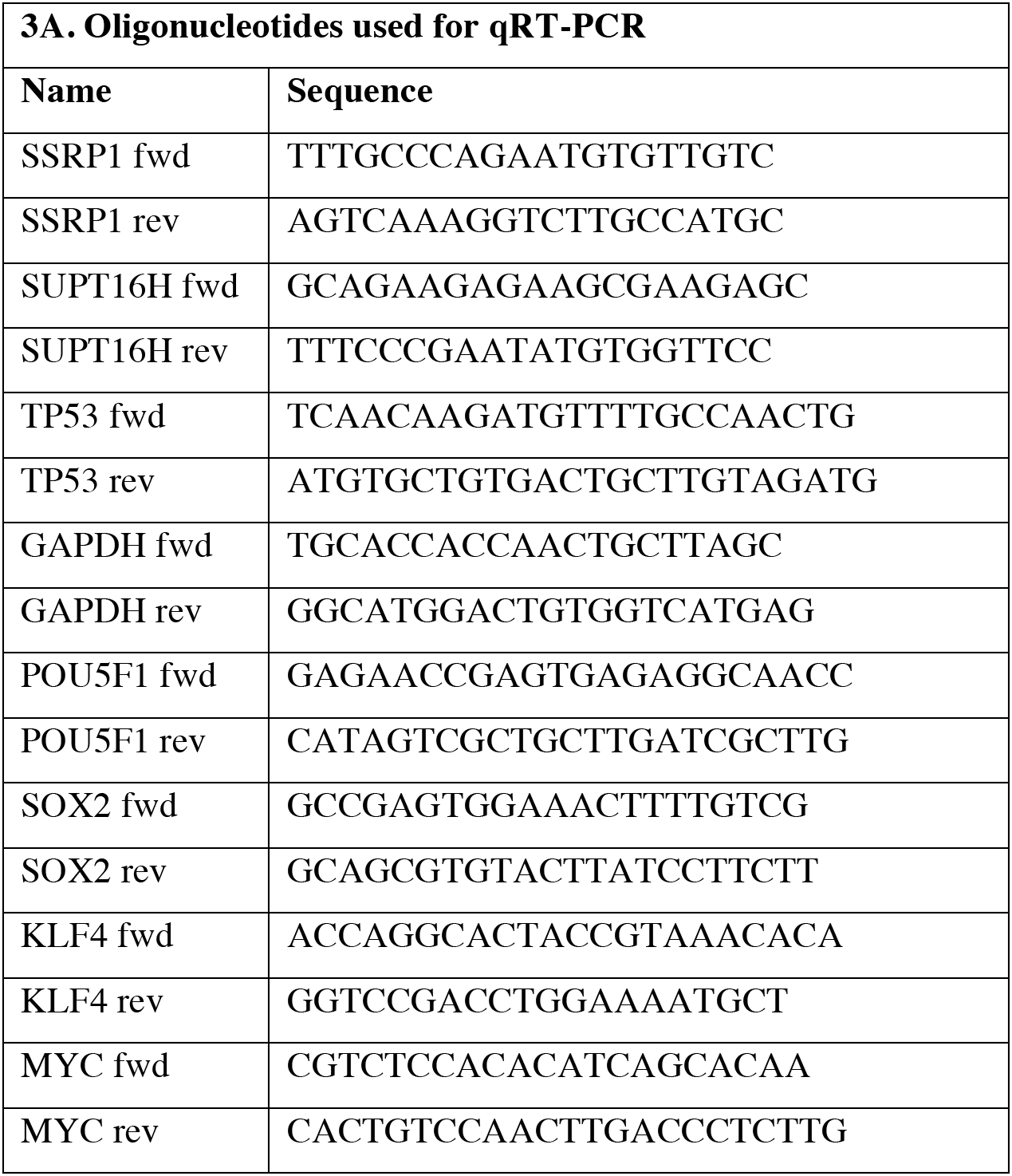
Oligonucleotides used for qRT-PCR and list of used siRNAs.

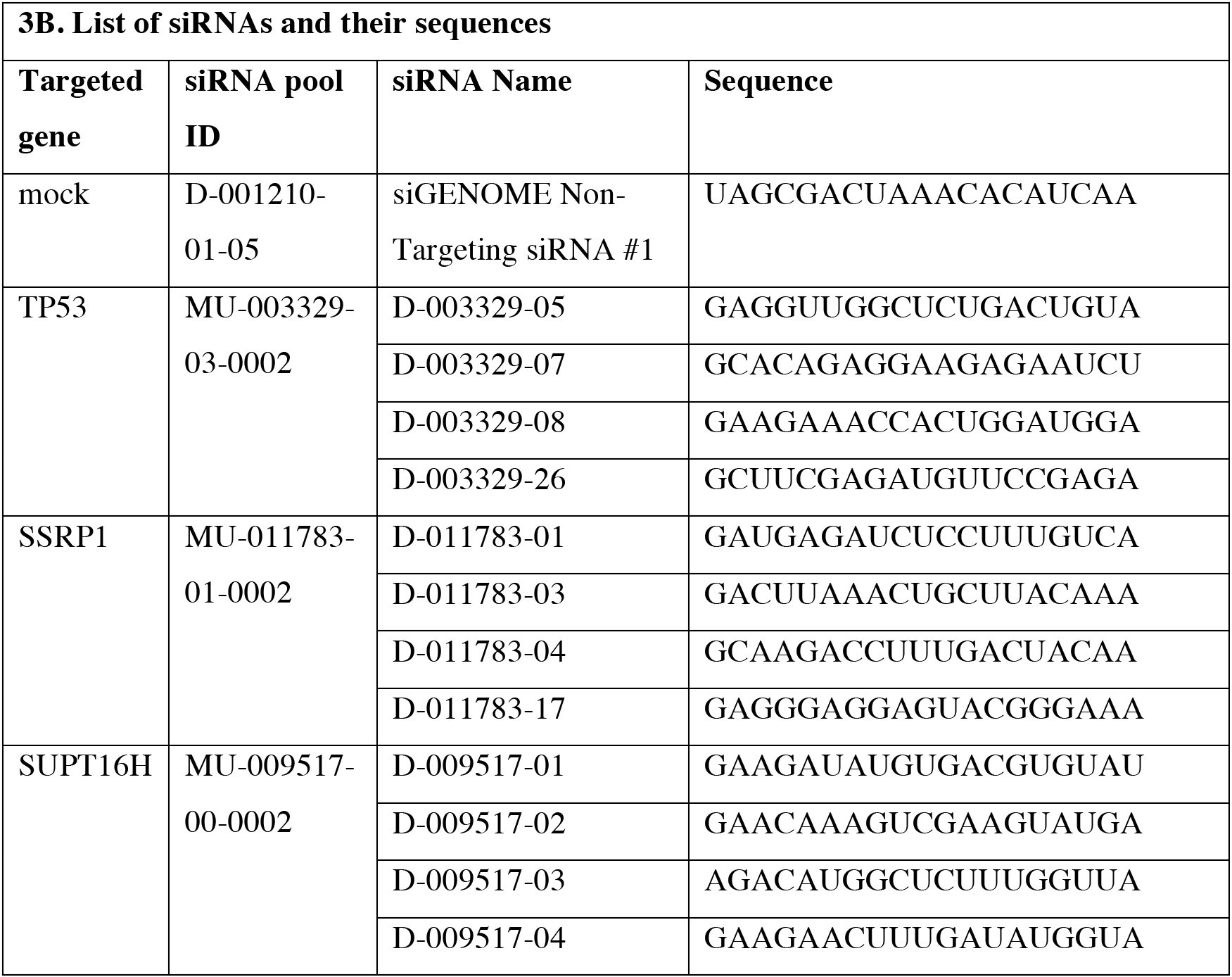
Oligonucleotides used for qRT-PCR and list of used siRNAs.

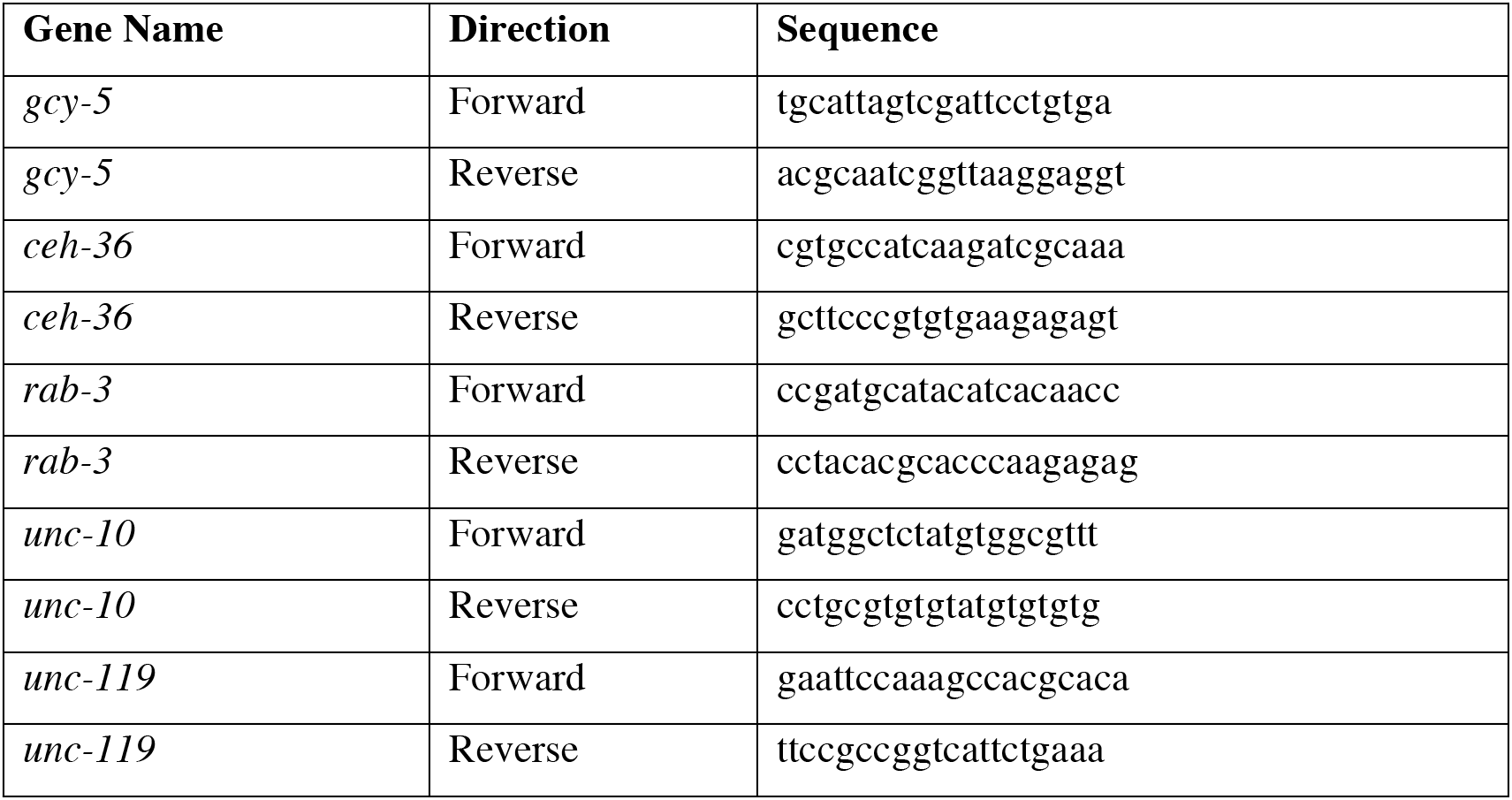
List of oligonucleotides used for qChIP.

### Nematode culture and RNAi

*C. elegans* strains were maintained on 0P50 bacteria at 20°C as described before(Brenner, 1974). All heat-shock and temperature-sensitive strains were kept at 15 °C. For RNAi, worms were grown on plates seeded with RNAi bacteria from Ahringer library (Source Bioscience). RNAi against *Renilla luciferase* (Rluc) was used as control. Reprogramming experiments were carried out either as P0 for *hmg-4* and *spt-16*, or F1 for *hmg-3* RNAi using a standard feeding protocol (Kamath et al., 2003). For P0 experiments, worms were synchronized by bleaching and L1 larvae were put on RNAi plates; for F1 RNAi, synchronized L1s were grown at 15°C on normal food until they reached L4 stage when they were transferred on RNAi plates. Worms on RNAi plates were grown at 15°C until most of the P0 or F1 progeny reached L4 stage. The plates were heat-shocked at 37°C for 30 min followed by an overnight incubation at 25°C as described before (Seelk et al., 2016; Tursun et al., 2011). Plates were screened for presence of ectopic GFP the following day under a dissecting scope. To induce the Glp phenotype in *glp-1(ar202)*, the animals were shifted to room temperature 8 hrs before the heat-shock. It is important to note that F1 RNAi against *spt-16* caused embryonic lethality thereby not allowing to test for *gcy-5::gfp* induction in the germline as seen upon *hmg-3* F1 RNAi. For double RNAi, bacteria were grown as saturated culture. The OD600 was measured to ensure that the bacteria were mixed in an appropriate 1:1 ratio and subsequently seeded on RNAi 6-well plates. The library screened for suppression and entire results are listed in Table S4. For time course experiments, screening for the presence of ectopic GFP was done 0, 4, 8, 12, 24, 36 and 48 hrs after heats-hock. For Aurora kinase B inhibitor experiments, ZM447439 (Selleckchem) was added to the NGM plates with a final concentration of 100 nM. The *C. elegans* lines used in this study are listed and described in detail in STAR methods.

### Generation of CRISPR alleles

CRISPR engineering was performed by microinjection using a PCR repair template and the *dpy-10* co-CRISPR approach as described before (Arribere et al., 2016; Paix et al., 2015). The injection mix contained a plasmid that drives expression of Cas9 (50ng/μl, a gift from John Calarco, Addgene #46168), one that drives expression of *dpy-10(cn64)* sgRNA (50ng/μl), a *dpy-10(cn64)* PAGE-purified 99mer single-stranded oligodeoxynucleotide (ssODN) HR template (50ng/μL; IDT), a plasmid expressing the sgRNA targeting the *hmg-3* locus (dBT620, 50ng/μl) and a PCR repair template to introduce the *3xHA* knock-in at the 3’ end of *hmg-3* (90ng/μl). To generate the sgRNA plasmids, annealed oligo pairs were ligated into BbsI-digested pJJR50 (a gift from Mike Boxem, Addgene#75026). To generate the PCR repair template, 3xHA tag was flanked with approximately 50bp homology arms on both 3’ and 5’ side for insertion at the 3’ end of the *hmg-3* locus Screening for successful knock-in events was done as described before (Arribere et al., 2016). Positive hits were homozygoused for the knock-in by singling and genotyping worms and the knock-in was confirmed by sanger sequencing.

Sequences of the oligonucleotides used for the generation of *hmg-3(bar24)* are reported in STAR methods.

### Antibody staining

For staining using anti-HMG-4/3 (rabbit polyclonal peptide antibody, 1:200) and anti-SPT16 (rabbit polyclonal peptide antibody, 1:200) antibodies, worms were resuspended 0,025 % glutaraldehyde and frozen using freeze-crack protocol (Duerr, 2006). Acetone/methanol fixation was used to prevent gonad extrusion. For anti-HA staining (anti-HA mono mouse antibody, Roche, at 1:1000 dilution) whole worms were fixed and permeabilized following a previously described method (Bettinger et al., 1996). In brief, after washing, worms were resuspended in RFB (160 mM KCl; 40 mM NaCl; 20 mM EGTA; 10 mM Spermidine) + 2% formaldehyde followed by three freeze-thaw cycles. After incubation for 30 min at 25°C, the sample was washed with TTE (100 mM Tris pH 7,4; 1 % Triton; 1mM EDTA) and incubated for 4 h at 37°C with shaking in TTE + 1% beta-Mercaptoethanol. The sample was washed in BO3 buffer (10 mM H_3_BO_3_; 10 mM NaOH; 2 % Triton) and further incubated for 15 min at 37°C with shaking in BO3 buffer + 10 mM DTT. After another wash with BO3, BO3 buffer + 0,3 % H_2_O_2_ was added and incubated for 15 min at 25°C. The sample was washed once more with BO3, blocked with 0,2 % gelatin + 0,25 % Triton in PBS and stained. Primary antibodies were diluted in PBS with 0,25 % Triton + 0,2 % gelatin, added to the fixed worms and incubated overnight at 4°C. After washing in PBS + 0,25 % Triton, secondary antibodies (Alexa Fluor dyes at 1:1500 dilution) were applied and incubated overnight at 4°C. Samples were washed in PBS + 0,25 % Triton and mounted on glass slides with DAPI-containing mounting medium (Dianove, #CR-3448).). For anti-P granule, ant-ELT-2 and anti-IFB-2 staining (anti-OIC1D4-s-P-granules, anti-455-2A4-s (ELT-2), anti-MH33-s – intermediate filament mono mouse antibody, repectively, Hybridoma bank, at 1:150 dilution) worms were dissected and processed as described before {Jones:1996fa}. The following antibodies were used: anti-OIC1D4-s (P-granules), anti-455-2A4-s (ELT-2), anti-MH33-s – intermediate filament (IFB-2), mono mouse antibodies, Hybridoma bank, at 1:150 dilution.

### Fosmid recombineering

Fosmid recombineering to create *barIs136* was performed as described in (Tursun et al., 2009). The recombineered fosmids were linearized and injected into hermaphrodites as complex arrays with 15 ng/ml of the linearized fosmid (WRM066C_F12 (Sarov et al., 2012)), 3 ng/ml of *myo-2^proī^::mCherry* used as a coinjection marker and 100 ng/ml of digested bacterial genomic DNA. Extrachromosomal lines acquired after injection were integrated into the genome using gamma irradiation.

### Single molecule fluorescent in situ hybridization (smFISH)

smFISH was performed using Custom Stellaris FISH probes, purchased from Biosearch Technologies and the staining was done according to the manufacturer’s protocol. Sequences of all the smFISH probes used are listed in STAR methods.

### Cell-Cycle Arrest by HU Treatment

Hydroxyurea (HU) treatment was carried out as previously described (Fox et al., 2011; Patel et al., 2012; Seelk et al., 2016). HU was added to seeded RNAi plates at a final concentration of 250 μM. L4 worms grown on RNAi plates were transferred to HU plates and incubated at room temperature for 5 hrs prior to heat-shock in order to induce CHE-1 expression. After overnight incubation, worms were assessed for GFP induction in the germline as described above.

### Cell culture and siRNA knockdown

Human secondary fibroblasts carrying a doxycycline (DOX)-inducible, polycistronic human OCT4/KLF4/c-MYC/SOX2 (OKMS) cassette (hiF-Tcells) (Cacchiarelli et al., 2015) were cultured in hiF medium (DMEM/F12 Gltuamax supplemented with 10% FBS, 1% NEAA, 0,1% beta-mercaptoethanol 100 U ml^-1^ penicillin, 100 μg ml^-1^ streptomycin and 16 ng/ml FGFbasic). hiF-T cells were passaged every 3 days, using a splitting ratio of 1:3 as described before(Cacchiarelli et al., 2015). For experiments, 80.000 hiF-T cells/ well of a 12-well plate treated with attachment factor were seeded and incubated overnight at 37°C. siRNA knockdown was performed the following day by a reverse transfection method using DharmaFECT 1 and 40 nM siRNA pool reagents purchased from Dharmacon according to the manufacturer’s instructions. Sequences of all the siRNA reagents used are described in STAR methods. To monitor knockdown efficiency, RNA was isolated using Qiagen RNeasy Plus Mini Kit 24h, 48h, 72h or 7 days after siRNA transfection. To control for OSKM expression after SSRP1 and SUPT16H depletion, RNA was isolated from siRNA transfected cells 48h after knockdown without prior Doxycycline treatment. cDNA was synthesized with oligo dT primers using the GoScript Reverse Transcriptase from Promega according to the manufacturer’s instructions. cDNA was used for qPCR with the Maxima SYBR Green qPCR Master Mix (2X) (Thermo Scientific) according to the manufacturer’s instructions on an ABI PRISM 7700 system (Applied Biosystems). The real-time PCR data analysis was done by using comparative C_T_ method (Schmittgen and Livak, 2008). Gene expression levels were calibrated to the housekeeping gene GAPDH and normalized to Rluc. Sequences of qRT-PCR primers used are described in STAR methods.

### Reprogramming experiments with hiF-T cells

Reprogramming experiments were performed by seeding hiF-T cells on irradiated mouse embryonic fibroblasts (MEF, GlobalStem) 24 h after siRNA transfection. Induction of OSKM by DOX supplementation (2μg/ml) was started the next day in hiF medium for the first 2 days and then in KSR medium (DMEM/F12 Gltuamax supplemented with 20% KSR, 1% NEAA, 0, 1% beta-mercaptoethanol 100 U ml^-1^ penicillin, 100 μg ml^-1^ streptomycin, 8 ng/ml FGFbasic and ROCK1 inhibitor (Y-27632-2HCl, Biozol Diagnostica, final concentration 1μM)) as described previously (Cacchiarelli et al., 2015).

### Reprogramming experiments with NHDF cells

Direct reprogramming experiments were performed by transducing Normal Human Dermal Fibroblasts (NHDF cells, Lonza) with RTTA, tetO-Ascl1, tetO-Brn2 and tetO-Myt1l (BAM) in a 1.2:1:1.5:2.5 ratio and supplemented with polybrene (c.f. 8 μg/ml) 24 h before siRNA transfection to deplete FACT. Induction of BAM by DOX supplementation (2 μg/ml) was started the next day in neuronal medium as described before (Vierbuchen et al., 2010)

### Phenotypic characterization of iPS cells

iPSC colonies were characterized after ~21 days of reprogramming. Alkaline phosphatase activity was measured using an enzymatic assay for alkaline phosphatase (Alkaline Phosphatase Staining Kit II, Stemgent) according to the manufacturer’s instructions. Number of colonies formed in each condition was counted based on SSEA-4 positive colonies. Immunohistochemistry with SSEA-4 Antibody (DyLight 488 conjugate, Thermo Scientific) was performed according to the manufacturer’s protocol at 1:500 dilution. Cells were fixed with 4%PFA in PBS and permeabilized with 0.1% Tween-PBS before antibody incubation. The pluripotency staining was performed using the Pluripotent Stem Cell 4-Marker Immunocytochemistry Kit (Thermo Fisher Scientific) in addition with the NANOG rabbit (Thermo, PA1-097) antibody.

### Pluripotency Teratoma Assay

The teratoma assay was done by EPO GmbH – Experimental Pharmacology & Oncology, Berlin, Germany. Briefly, in order to initiate the assay 1 x10^6^ cells were suspended in 50 ul of PBS and thereafter mixed with 50 ul of Matrigel (Corning). These cell suspension was than subcutaneously transplanted into a NOG-mice and the tumor growth was documented on a weekly basis. After reaching the size of 1,5cm^3^ the tumor was extracted and pathologically analyzed.

### Human ATAC-seq

For ATAC-seq of human fibroblasts, 50.000 cells were harvested 48 h after siRNA transfection and the cell pellet was resuspended in transposase reaction mix as described previously (Buenrostro et al., 2013). Briefly, cell pellet was resuspended in the transposase reaction mix (25 μL 2× TD buffer, 2.5 μL transposase) using Nextera DNA Library Preparation Kit (Illumina) and the transposition reaction was carried out for 60 min at 37°C. The samples were purified using Zymo DNA Clean & Concentrator kit. Following purification, library fragments were amplified using NEBNext PCR master mix and previously published PCR primers (Buenrostro et al., 2013). Libraries were amplified for a total of 12 to 14 cycles and sequenced using paired-end-sequencing length of 75 nucleotides using NextSeq 500/550 High Output v2 kit (Illumina).

### *C. elegans* nuclei isolation and ATAC-seq

For *C. elegans* ATAC-seq, L1 animals were synchronized by harvesting embryos using sodium hypochlorite treatment and grown on RNAi plates until L4 stage (P0 RNAi as described above). Synchronized L4 animals were washed 5 times in M9 buffer and collected on ice. Nuclei were isolated using a glass Dounce homogenizer with 50 strokes tight-fitting insert in buffer A (15 mM Tris-HCl pH7.5, 2 mM MgCl2, 340 mM sucrose, 0.2 mM spermine, 0.5 mM spermidine, 0.5 mM phenylmethanesulfonate [PMSF], 1mM DTT, 0.1% Trition X-100 and 0.25% NP-40 substitute) as described before (Ooi et al., 2010; Steiner et al., 2012). The debris were removed by spinning at 100×g for 5 min and nuclei were counted by Methylene blue staining. 100.000 nuclei per sample were pelleted by spinning at 1000×g for 10 min and proceeded immediately to transposition step of the ATACseq protocol as described above (Buenrostro et al., 2013). Libraries were amplified for a total of 10 to 18 cycles and sequenced using paired-end-sequencing length of 75 nucleotides using NextSeq 500/550 High Output v2 kit (Illumina).

### ATAC-seq analysis. *Pre-processing*

ATAC-seq reads were trimmed for adapters using flexbar v2.5 (-f i1.8 -u 10 -ae RIGHT -at 1.0) (Dodt et al., 2012) and mapped with bowtie2 v2.0.2 (Langmead and Salzberg, 2012) in default paired-end mode and restricting pair distances to 1500 (-X 1500 – no-discordant) to version hg19 of the human genome or ce10 of the worm genome followed by removal of multimappers. PCR duplicates were removed using Picard Tools MarkDuplicates v1.90 (http://broadinstitute.github.io/picard) and reads were converted to.bed format using bedtools bamToBed v2.23 (Quinlan and Hall, 2010). Pairs were filtered out if they mapped to the same strand, to different chromosomes, or if the 5’end coordinate of the – strand read was less than or equal to the 5’end coordinate of the + strand read. Mapped pairs were split into single reads and converted to a 38-bp fragment reflecting the theoretical minimal spacing required for a transposition event by Tn5 transpososome (Adey et al., 2010) using bedtools slop on the read 5’ends (-l 15 -r 22;) (Quinlan and Hall, 2010). Replicates were concatenated after confirming high concordance. *C. elegans* datasets were further filtered for reads from the rDNA loci as well as those mapping to regions corresponding to transgenic reporter constructs existing in the strains and corresponding to sequences used in the RNAi vectors. See Table S1 for basic statistics.

### Peak calling and differential analysis

Peaks were called on concatenated processed bed files using JAMM peakcaller v1.0.7.5 (-f38 -b 100 -e auto (human) and -e 1.75 (worm)) (Ibrahim et al., 2015). The resulting “all” (for meta analyses described below) or “filtered” peaks output by JAMM for each condition (control, SSRP1 knockdown, and SUPT16H knockdown for human, *rluc* and *hmg-3* knockdown, or rluc and *hmg-4, and spt-16* knockdown for worms) were concatenated and then merged with bedtools merge v2.23 (Quinlan and Hall, 2010). Merged “all” peaks were further filtered for a minimum width of 25-bp and used for meta analyses as described below. Merged “filtered” peaks for human (totaling 106,967) and 25-bp width-filtered merged “all” peaks for worm (totaling 17,024 for the *hmg-4/spt-16* experiment and 21,254 for the *hmg-3* experiment) were then counted for the number of processed reads that intersected them from each replicate of each experimental condition using bedtools coverage v2.23 (-counts) (Quinlan and Hall, 2010). Count tables were then normalized (method = “TMM”) and subjected to differential analysis (exactTest) using edgeR (Robinson et al., 2010). Peaks were called as differential with a q-value cutoff of 0.01 (adjust=“BH”).

### Meta analyses

Annotated transcript start sites from the gencode GTF v19 or the worm GTF were uniqued for gene IDs, collapsed for unique start coordinates, and intersected with merged “all” peaks described above using bedtools intersect v2.23 (Quinlan and Hall, 2010). Windows +/-1kb around this set of annotated start sites were then counted for their library-normalized coverage of replicate-concatenated processed ATAC-seq reads in each experimental condition, the resulting matrix averaged across positions and plotted using the ggplot2 R package (Wilkinson, 2011). Robust enhancer regions were downloaded from the FANTOM 5 database and intersected with the merged “all” peaks described above with bedtools intersect v2.23 (Quinlan and Hall, 2010). Worm ATAC-seq peaks greater than 500 bp from annotated transcription start sites as measured by bedtools closest v2.23 were taken as candidate enhancers. Windows +/-1kb around the midpoints of these human or worm candidate enhancers were then counted for their library-normalized coverage of replicate-concatenated processed ATAC-seq reads in each experimental condition, the resulting matrices averaged across positions and plotted using the ggplot2 R package (Wilkinson, 2011).

### Density scatter plots

Normalized log2 fold changes in each condition for peaks determined as differential (see above) in either SSSRP1 knockdown or SUPT16H knockdown or both for the human cells *(hmg-4* or *spt-16* knockdown for the worm data) were plotted against each other using ggplot2 geom_hex (bins=50) (Wilkinson, 2011).

### Peak annotations

“Filtered” human ATAC-seq peaks were annotated as promoter-proximal if they located +/-200bp from gencode v19 annotated start sites using bedtools closest (Quinlan and Hall, 2010), otherwise they were considered promoter distal.

### De novo motif generation

For the human motif generation, promoter distal peaks were summed for their edgeR normalized log2 fold changes for each condition, split into two groups according to summed changes greater (opening peaks) or less than (closing peaks) zero. The resulting summed changes were used as the ranking statistic for motif generation using the cERMIT program (parameters: seed_length=6, min_motif_length=6, max_motif_length=12, required_core_length=5, cluster_sim_threshold=0.8, hypegeom_p_val_cutoff=1.0e-30, num_random_runs=1000, max_gene_set_size_percentage_threshold=0.3, min_gene_set_size_percentage_threshold=0.01, degen_threshold=0.75, fraction_degen_pos_threshold=0.75, use_regression_scoring=no, bootstrap_motif_score=no, fast_mode=no; PMID: 20156354). The same analysis was also done with the log2 fold change ranks separate for each factor.

Worm motifs were generated the same way, except independent of peak annotation and log2 fold changes were used individually for each factor knockdown as the ranking statistic for cERMIT. Sequences were restricted to those with lengths <1000-bp and worm sequences for those >50-bp and <1000-bp prior to input into cERMIT. Sequence numbers put into cERMIT are as follows: human combined up 44068; human combined down 42720; SSRP1 up 45710, SSRP1 down 41087, SUPT16H up 42130, SUPT16H down 44667, *hmg-3* down 10653, *hmg-3* up 10486, *hmg-4* down 8721, *hmg-4* up 8221, *spt-16* up 8616, *spt-16* down 8326. cERMIT-generated motifs were converted to meme format and Tomtom (Gupta et al., 2007) was used in the default settings to match the generated motifs to the JASPAR CORE vertebrate 2016 database (Mathelier et al., 2016) for the human motifs or to a published database of protein binding microarray-generated motifs for *C. elegans* (Narasimhan et al., 2015).

### RNA-seq using *C. elegans*

For transcriptome analysis, RNA was isolated from HMG-4 and SPT-16-depleted animals using TRIzol (Life Technologies) and guanidinium thiocyanate-phenol-chloroform extraction. After adding chloroform to the sample containing TRIzol, phases were separated to an aquaous phase containing RNA, an interphase and an organic phase containing DNA and proteins. Guanidinium thiocyanate denatured proteins (including RNases) in the organic phase. RNA was purified from the aquaous phase using isopropanol. For reverse transcription GoScript Reverse Transcriptase (Promega) was used according to the manufacturers protocol. The preparation of libraries for whole-transcriptome sequencing was carried out using TruSeq RNA Library Prep Kit v2 (Illumina) according to the manufacturers instructions. Libraries were sequenced using single end sequencing length of a 100 nucleotides on a HiSeq4000 machine (Illumina).

## Analysis of RNA-seq

Multiplexed Illumina sequencing data was demultiplexed using Illumina bcl2fastq converter (version v2.17.1.14). Raw reads in fastq format were processed to get rid of low quality bases and possible adapter contamination using Trimmomatic (version 0.33) (Bolger et al., 2014) (settings: ILLUMINACLIP:TruSeq.fa:2:25:6 LEADING:3 TRAILING:3 SLIDINGWINDOW:4:15 MINLEN:36). The filtered reads were aligned to the *C.elegans* genome (ce10 genome build – WormBase WS220 released in October, 2010) using the splice-aware short read aligner STAR (version 2.4.2a) with the default settings (Dobin et al., 2012)except for setting “ — outFilterMultimapNmax” argument to

1. The expression level of each gene was quantified using R/Bioconductor package quasR (Gaidatzis et al., 2014) using genome annotation data in GTF file format from the Ensembl database (version 70) (Yates et al., 2016) Differential expression analysis of the quantified expression levels of genes between different samples was done using the R/Bioconductor package DESeq2 (Love et al., 2014) Up/down regulated genes are detected based on the differential expression criteria of adjusted p-value of at least 0.1 and at least two-fold increase/decrease in expression levels in relation to the control samples

### Data availability

The following link has been created to allow review of record GSE98758 while it remains in private status: upon request.

